# Programmable Biochemical Logic Through Repurposing Natural bZIP Protein Interaction Networks

**DOI:** 10.64898/2026.07.29.741539

**Authors:** Arturo Orozco-Estrada, Daniela Flores-Nuño, Gerardo Mendizabal-Ruiz, Ernesto Borrayo, J. Alejandro Morales

## Abstract

Protein dimerization networks can implement biochemical logic, but their practical realization has largely relied on designing or optimizing protein–protein interaction affinities, creating a major barrier to experimental implementation. Here, we ask whether computation can instead be extracted from interaction networks already generated by evolution. We repurpose experimentally characterized basic leucine zipper (bZIP) interactomes from *Nematostella vectensis*, *Ciona intestinalis*, and *Homo sapiens* as programmable biochemical logic substrates while keeping all measured binding affinities fixed. Using mass-action equilibrium modeling coupled with genetic optimization, we identify accessory monomer concentrations and output dimers that reconfigure these natural networks to perform Boolean operations. Unengineered bZIP clusters computed up to 15 of the 16 possible two-input Boolean functions, including nonlinearly separable XOR and XNOR operations, and up to 13 functions within a 17-gate three-input benchmark. Computational versatility depended on network architecture and input selection, revealing design constraints inherent to natural interactomes. Stochastic perturbation analysis further showed that many optimized configurations preserve logic-state separation under substantial concentration variability. Finally, a nondimensional scaling framework provides a route for translating simulated concentration regimes toward cell-free implementation. Together, these results demonstrate computationally that natural protein interaction networks contain latent, evolutionarily encoded computational capacity that can be accessed through concentration tuning rather than *de novo* protein engineering.

**Graphical abstract:** 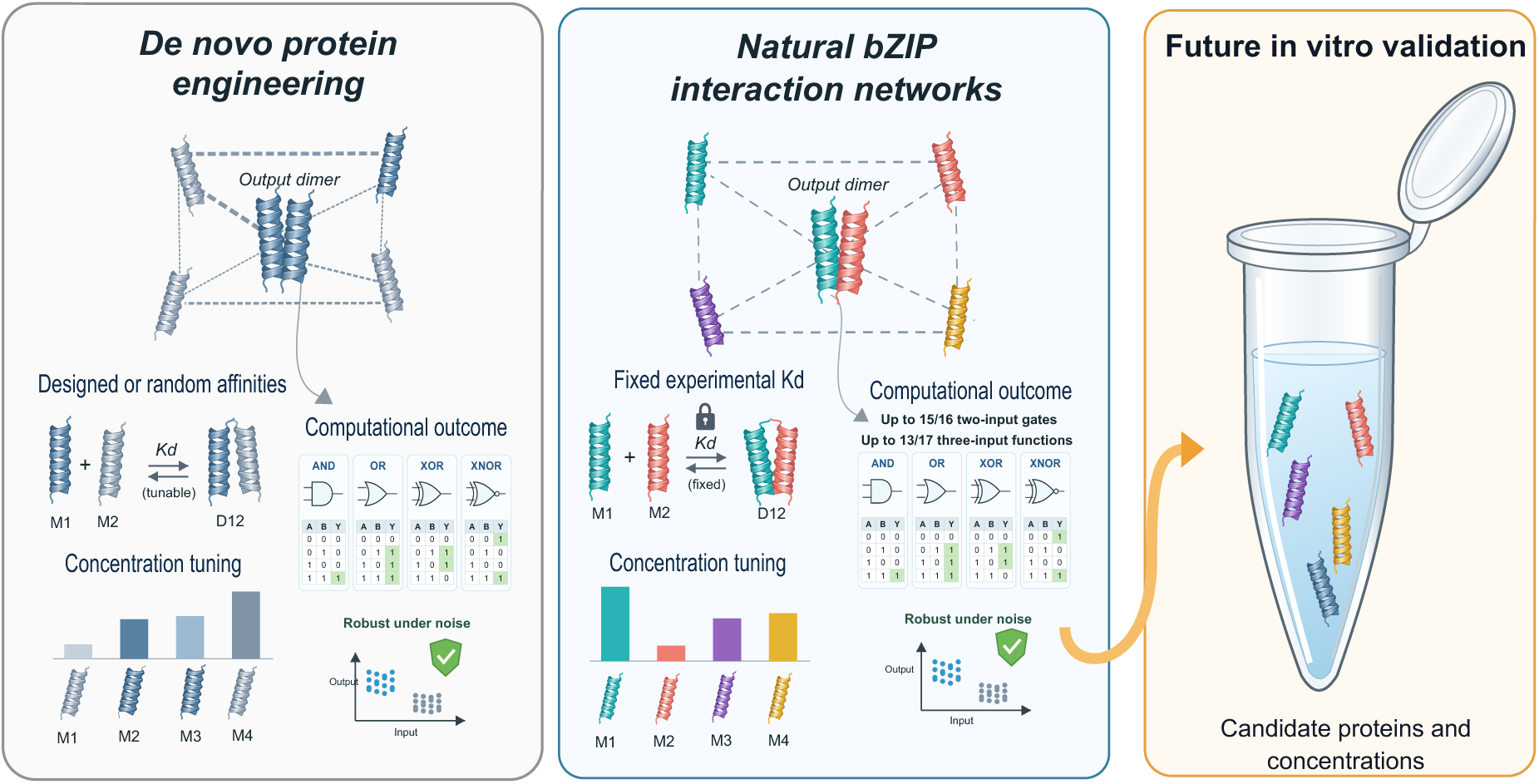

## 1. Introduction

Modern computing faces an escalating energy crisis driven by the unsustainable power demands of silicon-based artificial neural network architectures^1^. Biological computing offers a transformative alternative, processing complex information through molecular interactions under chemical equilibrium near fundamental thermodynamic limits of energy efficiency^2^. Beyond energy efficiency, this molecular approach also offers opportunities for molecular biosensing and cell-based biological processors^3^. At the diagnostic and therapeutic frontier, however, conventional biosensors remain severely limited: they typically monitor single biomarkers, lack multi-input decision-making capacity, and exhibit slow response dynamics^4^. Within synthetic biology, logic processing has traditionally relied on gene regulatory circuits^5,6^. Yet, these genetic networks are fundamentally constrained by transcription-translation-dependent response times of hours^7^ and impose heavy metabolic burden and cellular toxicity when scaled to evaluate multi-signal inputs^8^. To enable rapid, energy-efficient molecular decision-making, competitive protein dimerization networks have emerged as a powerful alternative^9^. These networks rely on direct protein–protein interactions within a single biochemical layer to execute multi-input Boolean operations within seconds^10^, with substantially lower metabolic costs^8^.

In dimerization networks, upstream molecular signals dictate the concentrations of specific protein monomers (inputs), which competitively associate to form biologically active homodimers or heterodimers (outputs)^9^. This biochemical computation is prevalent in nature and serves as a fundamental regulatory motif in biological families such as nuclear receptors, leucine zippers, and helix-loop-helix proteins^11–13^. Recent theoretical frameworks have demonstrated that competitive dimerization networks, modeled under mass-action equilibrium (**Section 5.1**), are mathematically capable of computing complex Boolean functions^9,14,15^. A network’s computational behavior is dictated by two key variables: the pairwise binding affinities (*K_ij_*) of the monomers and the total concentrations of accessory monomers (*M_i_*). By optimizing these parameters, networks achieve high computational versatility, defined as the number of distinct logic gates a single interactome can execute purely by modulating accessory species abundances^9,14^.

Despite their theoretical power, the physical implementation of these optimized networks *in vitro* or *in vivo* faces a challenging bioengineering bottleneck: the requirement for custom proteins with strictly predefined equilibrium constants. Designing *de novo* proteins based exclusively on targeted interaction affinities remains a difficult structural challenge^16,17^. Although previous theoretical studies circumvented this by assigning random binding affinities^9^, such unconstrained parameter spaces fail to reflect the physical, structural, and evolutionary constraints that shape real biological protein families^9,18,19^.

To overcome the limitations of *de novo* protein engineering^16,17^, an alternative paradigm is to repurpose natural protein interactomes with pre-characterized binding profiles. The basic leucine zipper (bZIP) transcription factor family, defined by a basic DNA-binding domain and a leucine zipper region mediating homo- and heterodimerization^20,21^, is an ideal substrate as its comprehensive interactome and pairwise equilibrium constants have been experimentally mapped^18^. By fixing their experimentally measured *K_ij_* values, we can achieve multi-input biochemical computation solely by modulating accessory monomer concentrations ([*M_i_*]). This approach shifts the bioengineering bottleneck from complicated structural protein design to manageable chemical concentration tuning.

The latent computational capacity of unengineered protein networks provides a foundation for the design of robust, energy-efficient biocomputing units that can execute complex, multi-input logic.

Such programmable molecular processors are essential for the future development of rapid cell-free diagnostic platforms capable of discriminating intricate pathogen signatures and intelligent cell-based therapies that process multi-signal disease microenvironments in real time.

Here, we establish an integrated *in silico* framework to systematically evaluate the computational versatility and robustness of unengineered natural bZIP interactomes across *Nematostella vectensis*, *Ciona intestinalis*, and *Homo sapiens* derived from experimental binding datasets mapped by Reinke *et al.*^18^. First, we apply weighted Louvain community detection to isolate densely connected topological clusters and reduce system complexity and signal dissipation (**Section 2.1**). Next, we model mass-action chemical equilibria coupled with a genetic algorithm to identify the accessory monomer concentration profiles ([*M_i_*]) required to execute two-input (**sections 2.2 and 2.3**) and three-input (**section 2.4**) Boolean logic operations while holding experimental binding affinities (*K_ij_*) strictly constant. We then evaluate the functional resilience of these optimized logic gates under stochastic, Gamma-distributed protein expression noise (**Section 2.5**). Finally, we formulate a non-dimensional thermodynamic scaling framework to bridge nanomolar simulation parameters with higher-concentration *in vitro* cell-free diagnostic assays (**section 3**). Together, our findings demonstrate that natural protein interaction networks harbor rich, evolutionarily tuned biocomputing capacity, which provides a viable foundation for protein-based biocomputing without the need for *de novo* protein engineering.

## 2. Results and Discussion

Biomolecular computation operating via competitive protein dimerization bridges the gap between physical chemistry and biological systems through the integration of mass-action equilibrium thermodynamics with the functional diversity of evolved interactomes. Harnessing these interactions as processing modules for future biosensors and cell-based therapies requires establishing which logical functions they can support, how their behavior can be reconfigured, and whether their outputs remain distinguishable when component concentrations vary. We address these questions using experimentally characterized natural bZIP clusters.

This section details the systematic characterization of natural bZIP interactomes for molecular logic computation. We first present the structural characterization of the selected bZIP clusters. Subsequently, we evaluate their computational versatility across two-input and three-input Boolean operations optimized via genetic algorithms. To conclude, we test the functional resilience of the optimized networks against biologically realistic expression noise and evaluate the concentration scaling requirements for physical implementation.

### 2.1. Community detection and cluster extraction

A practical barrier to implementing natural protein networks as molecular processors is the complexity of reconstructing complete interactomes. Producing and controlling numerous molecular species increases experimental demands^22^ and, in cellular implementations, can impose additional burden on cellular expression resources^23^. Extensive background crosstalk can also redistribute components through competing interactions, which complicates the relationship between designated inputs and outputs^9,24^. Repurposing natural interactions therefore requires determining whether smaller protein assemblies can retain the competitive interactions needed for computation. Community extraction addresses this architectural challenge by defining restricted sets of experimentally characterized components, allowing us to investigate molecular logic without reconstructing entire interactomes.

To reduce system complexity, we partitioned the species-specific bZIP networks of *Homo sapiens, Nematostella vectensis,* and *Ciona intestinalis* into smaller communities via the weighted Louvain algorithm^25^, with experimental association constants (*K_a_*) as edge weights (**Section 5.3**). By maximizing modularity, this procedure identifies groups with strong internal connectivity that provide candidate substrates for subsequent logic optimization. The complete partitioning results and network visualizations are provided in **Supplementary Note S1**.

For clarity throughout the remainder of this manuscript, we establish the following terminology. We define a complete network as the full bZIP protein family of a given organism, a cluster as a subset of proteins extracted from a complete network, and an individual network as a cluster with specific accessory monomer concentrations that is capable of computing at least one logical gate.

We filtered the partitioned communities based on their size and computational potential and selected a total of 13 clusters (**section 5.3**). Their structural properties and internal connectivity are consolidated in **Table 1**. The topological features and species-specific interaction profiles of these selected modules are detailed below for each organism.

**Table 1.**
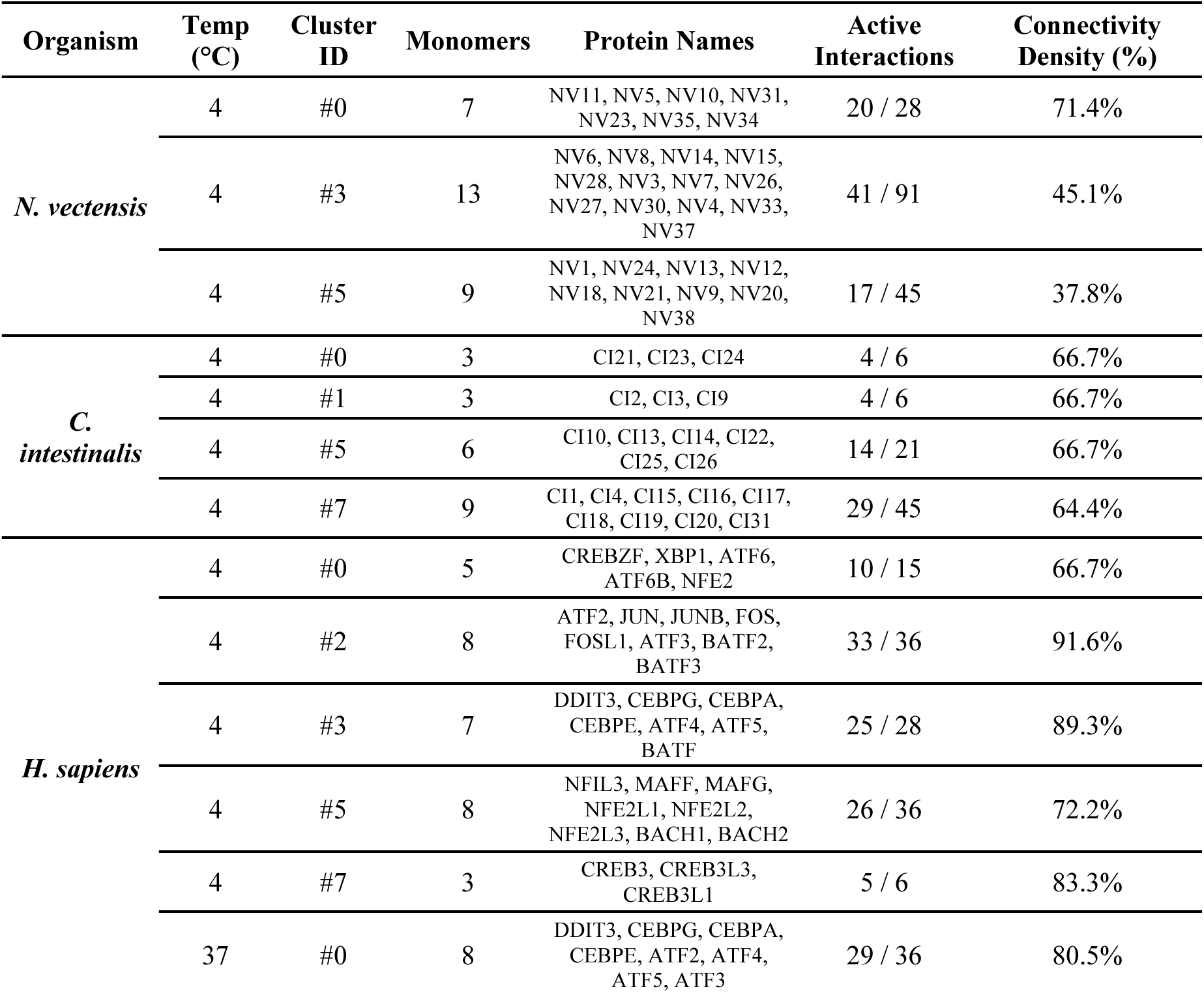
Properties of the bZIP clusters selected for computational analysis. For each cluster, the organism, experimental temperature, cluster identifier, number of monomers, monomer name, number of active protein-protein interactions out of the possible number of theoretical interactions, and connectivity density are reported. Monomer names listed in the *Protein Names* column are ordered sequentially to match the zero-indexed numerical tags (0, 1, 2, …) displayed in the interactome network figures (**fig 1 and figures in supplementary material S1**).

| Organism | Temp (°C) | Cluster ID | Monomers | Protein Names | Active Interactions | Connectivity Density (%) |
|---|---|---|---|---|---|---|
| ***N. vectensis*** | 4 | #0 | 7 | NV11, NV5, NV10, NV31, NV23, NV35, NV34 | 20 / 28 | 71.4% |
|  | 4 | #3 | 13 | NV6, NV8, NV14, NV15, NV28, NV3, NV7, NV26, NV27, NV30, NV4, NV33, NV37 | 41 / 91 | 45.1% |
|  | 4 | #5 | 9 | NV1, NV24, NV13, NV12, NV18, NV21, NV9, NV20, NV38 | 17 / 45 | 37.8% |
| ***C. intestinalis*** | 4 | #0 | 3 | CI21, CI23, CI24 | 4 / 6 | 66.7% |
|  | 4 | #1 | 3 | CI2, CI3, CI9 | 4 / 6 | 66.7% |
|  | 4 | #5 | 6 | CI10, CI13, CI14, CI22, CI25, CI26 | 14 / 21 | 66.7% |
|  | 4 | #7 | 9 | CI1, CI4, CI15, CI16, CI17, CI18, CI19, CI20, CI31 | 29 / 45 | 64.4% |
| ***H. sapiens*** | 4 | #0 | 5 | CREBZF, XBP1, ATF6, ATF6B, NFE2 | 10 / 15 | 66.7% |
|  | 4 | #2 | 8 | ATF2, JUN, JUNB, FOS, FOSL1, ATF3, BATF2, BATF3 | 33 / 36 | 91.6% |
|  | 4 | #3 | 7 | DDIT3, CEBPG, CEBPA, CEBPE, ATF4, ATF5, BATF | 25 / 28 | 89.3% |
|  | 4 | #5 | 8 | NFIL3, MAFF, MAFG, NFE2L1, NFE2L2, NFE2L3, BACH1, BACH2 | 26 / 36 | 72.2% |
|  | 4 | #7 | 3 | CREB3, CREB3L3, CREB3L1 | 5 / 6 | 83.3% |
|  | 37 | #0 | 8 | DDIT3, CEBPG, CEBPA, CEBPE, ATF2, ATF4, ATF5, ATF3 | 29 / 36 | 80.5% |

#### 2.1.1. Nematostella vectensis Clusters (4°C)

The *N. vectensis* network was partitioned into 8 distinct clusters. We selected three for optimization: **Cluster #0** (7 monomers), identified as the densest community in this organism with 71.4% connectivity density (20 active interactions out of 28 possible pairs) (**Fig. 1A**); **Cluster #3** (13 monomers), which provides the largest absolute competitive background with 41 active interactions (**Fig. S4**); and **Cluster #5** (9 monomers), which exhibits a more moderate density of 17 active interactions (**Fig. S5**).

**Fig. 1.**
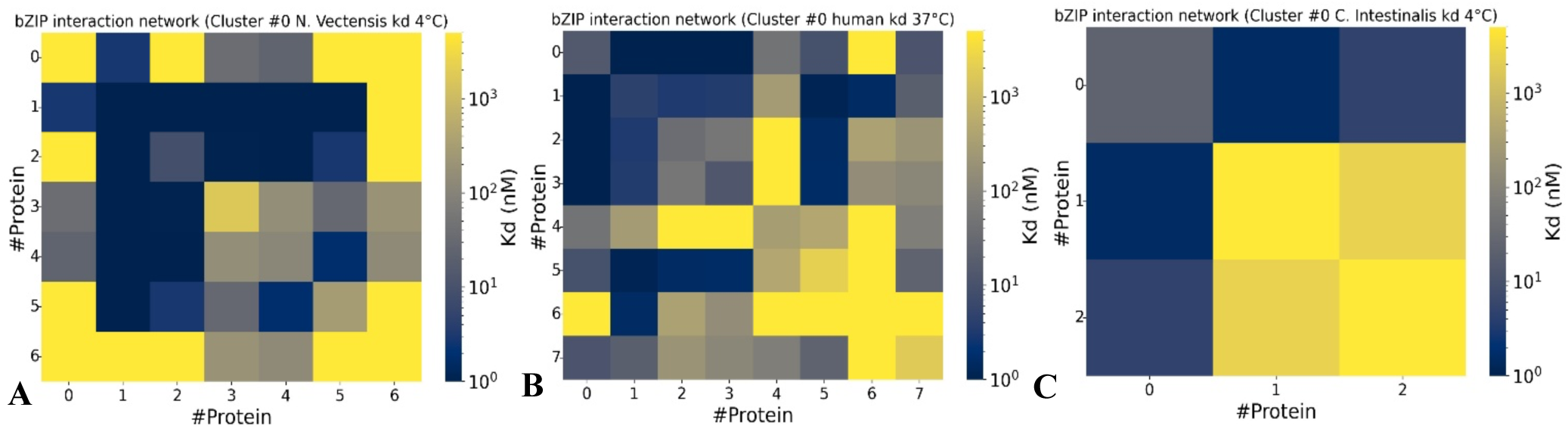
Representative heatmaps of bZIP interaction networks used as computational substrates, depicting the experimentally determined pairwise dissociation constants (*K_d_*) for representative clusters from **A)** *N. vectensi*s (Cluster #0, 4°C), **B)** *H. sapiens* (Cluster #0, 37°C), and **C)** *C. intestinalis* (Cluster #0, 4°C). Color intensity indicates the magnitude of the dissociation constant for each protein-protein interaction within each cluster; stronger interactions are presented in blue.

#### 2.1.2. *Ciona intestinalis* Clusters (4°C)

The *C. intestinalis* network yielded 8 highly isolated clusters with minimal inter-cluster crosstalk. We selected four structurally viable substrates: **Cluster #0** (**Fig. 1C**) and **Cluster #1** (3 monomers each), representing minimal, exceptionally dense, and tightly coupled three-protein networks (**Fig. S9**); **Cluster #5** (6 monomers) with a cohesive internal topology (14 interactions) (**Fig. S10**); and **Cluster #7** (9 monomers), which mirrors the density of the smaller communities with 29 active internal connections (**Fig. S11**).

#### 2.1.3. *Homo sapiens* Clusters (4°C and 37°C)

At 4°C, the human network was partitioned into 8 clusters, from which we selected five: **Cluster #0** (5 monomers, 10 interactions) (**Fig. S16**); **Cluster #2** (8 monomers), which is highly saturated with 91.6% connectivity density (33 out of 36 interactions) (**Fig. S17**); **Cluster #3** (7 monomers, 25 interactions) (**Fig. S18**); **Cluster #5** (8 monomers, 26 interactions), represents the least dense community (**Fig. S19**); and **Cluster #7** (3 monomers, 5 interactions) exhibits a high density of 5 interactions out of 6 possible pairs (**Fig. S20**).

At physiological 37°C, elevated thermal energy significantly constrains protein-protein affinities^18^ and reduces the total stable dimers across the human network (**section 5.3**). Modularity optimization at 37°C still yields 8 communities, but with a marked reduction in inter-cluster crosstalk. Under these conditions, we isolated Human **Cluster #0** (37°C) (8 monomers, 29 interactions), which corresponds topologically to **Cluster #3** at 4°C, to evaluate biocomputing performance under physiological parameters (**Fig. 1B**).

#### 2.1.4. Discussions of Structural Topology and Computational Latency

The selected bZIP clusters provide defined assemblies of naturally interacting proteins, which bring the proposed use of natural interactomes closer to experimentally testable configurations. By restricting subsequent optimization to these smaller subsets, we reduce the number of molecular species whose production and concentrations would need to be controlled during reconstruction. This directly addresses an implementation difficulty associated with complete interactomes: the coordination of numerous components and their competing interactions. In cellular implementations, using fewer expressed components may also help limit resource demands, although this potential benefit requires experimental assessment^23^. The structural differences among the selected clusters therefore provide a basis for examining how distinct natural interaction architectures support computation within more restricted sets of components.

A cross-species comparison of these topologies reveals a clear evolutionary trade-off in network structure^18^. *Homo sapiens* and *N. vectensis* present highly integrated networks with high edge densities and substantial cross-talk between communities. Conversely, *C. intestinalis* presents a highly compartmentalized architecture where functional groups are naturally insulated from external perturbations (**Supplementary note S1-Figs. S2, S7, S13**).

**Fig. 2.**
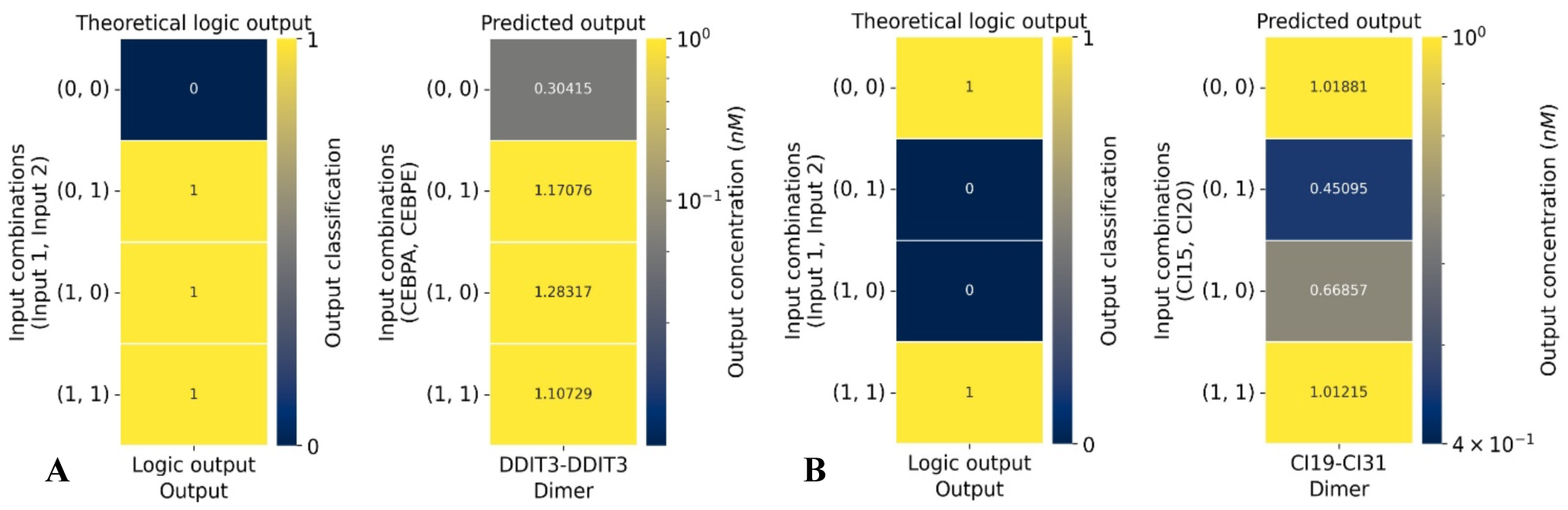
Representative two-input logic-gate responses produced by optimized bZIP individual networks. In each example, the theoretical truth table is shown on the left, while the predicted equilibrium concentration of the selected output dimer is shown on the right. **A)** The first case shows an OR gate computed by H. *sapiens* cluster #3 at 4°C. **B)** The second case shows an XNOR gate computed by *C. intestinalis* cluster #7 at 4°C. Each row represents a distinct input combination, and color intensity in the predicted-output panels denotes the simulated output-dimer concentration.

From a biocomputing perspective, these structural differences offer distinct advantages. The insulated clusters of *C. intestinalis* provide ideal substrates for standalone, orthogonal logic units that can be implemented with minimal risk of unintended interaction with neighboring cellular components. On the other hand, the highly interconnected frameworks of *H. sapiens* and *N. vectensis* offer a vastly superior competitive background. The extensive cross-talk and high local densities within these clusters increase the complexity of the molecular equilibrium.

Crucially, the uniform absence of centralized hubs across all three organisms confirms that bZIP networks rely on distributed, scale-free competitive dynamics rather than a few master-regulator proteins^26^. This distributed architecture represents a major advantage for synthetic biology design: it implies that the computational capacity of the network is a systemic property of the community, although their robustness may still depend on specific key components^27^.

Taken together, the topological characterization and community extraction identify smaller bZIP assemblies. These results address the complexity of working with complete interactomes by narrowing the design space to specific protein subsets. The resulting set of architectures provides a structural basis for subsequent experimental assays, it also specifies which proteins would need to be produced and controlled, turning natural interaction datasets into concrete starting points for molecular circuit development.

### 2.2. Two-Input Logic Gates and Computational Versatility

Previous theoretical studies have demonstrated Boolean computation in networks with fixed interaction affinities, but achieving broad computational versatility in random-affinity networks has required relatively large assemblies^9^. Here, we propose that similar versatility can be obtained within smaller networks whose interactions are constrained by experimentally characterized binding affinities. We evaluate two-input Boolean logic in selected natural bZIP clusters. By optimizing accessory concentrations and selecting output dimers, we test how much computational versatility these defined architectures can support.

To evaluate whether natural protein interactomes can execute digital biocomputing, we modeled steady-state competitive dimerization across the 12 extracted bZIP clusters. To simulate logical computation under chemical equilibrium, we coupled mass-action thermodynamic calculations using the Equilibrium Toolkit (EQTK)^28,29^ with a continuous genetic algorithm that optimizes accessory monomer concentrations ([*M_i_*]) while experimental binding affinities (*K_ij_*) remain strictly fixed (**Sections 5.4–5.5**). Across the distinct topological structures of *N. vectensis*, *C. intestinalis*, and *H. sapiens*, this optimization framework identified a cumulative total of 135 functional individual networks capable of executing two-input Boolean operations under strict, non-overlapping logic-state separation criteria. As summarized in **Table 2**, natural bZIP clusters differ substantially in their computational versatility, defined as the fraction of the 16 possible two-input logic functions successfully computed.

**Table 2.** Computational versatility of selected bZIP clusters evaluated across all 16 two-input Boolean functions. For each cluster, the number of monomers, number of dimers/interactions, successfully computed logic gates, total evaluated Boolean functions, and final versatility score are reported.

| Organism | Cluster | Number of monomers | Number of possible dimers / interactions | Number of active dimers / interactions | Connection density | Computed logic gates | Theoretical two input logic gates | Versatility |
|---|---|---|---|---|---|---|---|---|
| ***NV*** | #0 | 7 | 28 | 20 | 0.71 | 14 | 16 | 0.8750 |
| ***NV*** | #3 | 13 | 91 | 41 | 0.45 | 15 | 16 | 0.9375 |
| ***NV*** | #5 | 9 | 45 | 17 | 0.38 | 12 | 16 | 0.7500 |
| ***CI*** | #0 | 3 | 6 | 4 | 0.67 | 5 | 16 | 0.3125 |
| ***CI*** | #1 | 3 | 6 | 4 | 0.67 | 7 | 16 | 0.4375 |
| ***CI*** | #5 | 6 | 21 | 14 | 0.67 | 12 | 16 | 0.7500 |
| ***CI*** | #7 | 9 | 45 | 29 | 0.64 | 15 | 16 | 0.9375 |
| ***HS*** | #0 | 5 | 15 | 10 | 0.67 | 11 | 16 | 0.6875 |
| ***HS*** | #2 | 8 | 36 | 33 | 0.92 | 14 | 16 | 0.8750 |
| ***HS*** | #3 | 7 | 28 | 25 | 0.89 | 14 | 16 | 0.8750 |
| ***HS*** | #5 | 8 | 36 | 26 | 0.72 | 11 | 16 | 0.6875 |
| ***HS*** | #7 | 3 | 6 | 5 | 0.83 | 5 | 16 | 0.3125 |
| ***HS 37°*** | #0 | 8 | 36 | 29 | 0.81 | 12 | 16 | 0.7500 |

The most versatile substrates were *C. intestinalis* cluster #7 and *N. vectensis* cluster #3, each computing 15 of the 16 possible gates (93.75%). High versatility was a cross-species trait, as multiple communities successfully computed 14 functions. Conversely, minimal network architectures established lower performance boundaries, as the smallest three-monomer communities were restricted to computing between 5 and 7 logic gates. These results indicate that natural bZIP clusters can support broad logical behavior, but that computational versatility varies with the size and connectivity of the cluster.

A second summary of the results is shown in **Supplementary note S2**-**Table S5**, where the specific gates computed by each cluster are reported across the complete set of 16 two-input Boolean functions.

Across all evaluated clusters, linearly separable operations (e.g., FALSE, AND, BUFFER A) were universally computed. Crucially, multiple natural clusters successfully implemented non-linearly separable functions (XOR and XNOR), which require highly coordinated competitive backgrounds where intermediate heterodimers dynamically sequester free monomers to achieve non-monotonic input-output relationships^9,30^. The successful implementation of non-linearly separable functions demonstrates that these natural clusters do not merely generate proportional, additive responses to separate inputs.

To illustrate this molecular processing, **Fig. 2** highlights the computation of two representative individual networks. In the first panel (**Fig. 2A**), a human **cluster #3** (at 4°C) maps a robust OR gate. In the second panel (**Fig. 2B**), *C. intestinalis* **cluster #7** executes a complex XNOR operation, which demonstrates that appropriate accessory monomer balancing can exploit competitive dimerization to produce advanced combinatorial logic from a single molecular layer.

Correlation analysis between cluster architecture and computational capacity revealed a positive trend: computational versatility scales linearly with both the number of monomers (R^2^ ≈ 0.70) and the total number of active interactions (R^2^ ≈ 0.73) (**Fig. 3**). These high coefficients of determination confirm that network scale and connectivity density are primary structural drivers of biocomputing capacity in natural bZIP clusters.

**Fig. 3.**
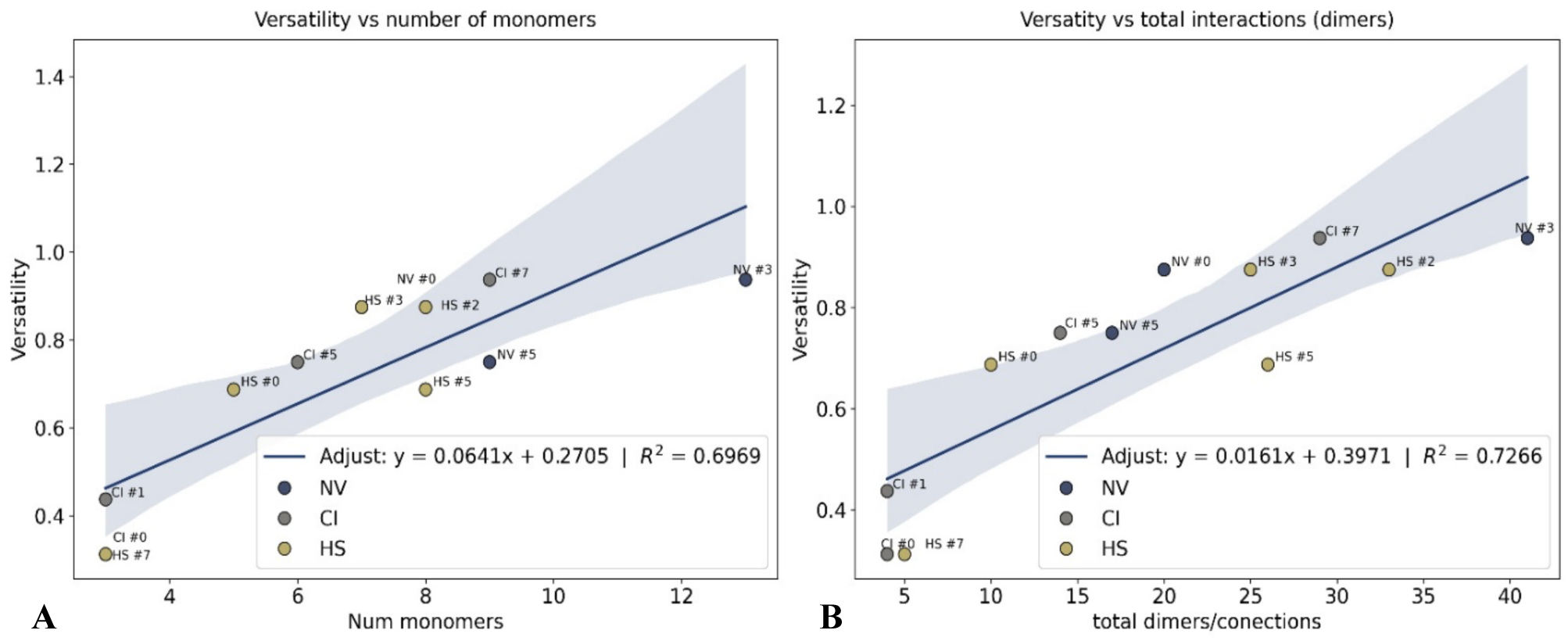
Association between bZIP cluster architecture and computational versatility. **A)** Versatility plotted against the number of monomers in each selected cluster. **B)** Versatility plotted against the total number of active dimers/interactions. Each point represents a selected bZIP cluster labeled by organism and cluster ID, with colors indicating the organism of origin. Solid lines indicate linear regression fits with corresponding coefficients of determination (R^2^), and shaded regions represent the 95% confidence intervals.

#### 2.2.1 Discussions of Two-Input Logic Computation and Natural Cluster Versatility / Evolutionary optimization vs. de novo design

Mathematically, a greater number of molecular species expands the dimensionality of the system’s mass-action equations, providing the genetic algorithm with a flexible parametric landscape to discover non-overlapping equilibrium states. Consequently, as a cluster grows in size and connection density, its internal competitive background becomes increasingly sophisticated, which significantly enhances its capacity to evaluate complex logic^30^.

However, comparing our natural bZIP models against previous random-affinity frameworks highlights a remarkable structural efficiency. In synthetic optimization studies^9^, networks with fixed random K values required a minimum of 20 monomers to successfully evaluate a benchmark suite of 10 two-input logic functions. In contrast, natural bZIP clusters achieved up to 15 out of 16 logic gates using clusters composed of only 9 and 13 monomers (*C. intestinalis* #7 and *N. vectensis* #3, respectively). Furthermore, comparing our natural interaction substrates against fully co-optimized synthetic frameworks further underscores the computational potency of natural interactomes. Previous theoretical work by Parres-Gold *et al.*^9^ demonstrated that co-optimizing both pairwise association constants (*K_a_*) and accessory monomer concentrations allowed synthetic networks of 3 to 6 monomers to compute a broad array of two-input functions, including non-linear operations such as XOR and XNOR. Based on our previous minimal network framework^14^, we conducted an additional assay where both *K_a_* values and accessory concentrations were co-optimized (**Supplementary Note S3**). In this unconstrained minimal setting, 3-monomer synthetic individual networks computed 14 out of the 16 two-input logic gates (all except XOR and OR), while the remaining XOR and OR gates were successfully resolved upon expanding the networks to 4 monomers (**Supplementary Note S3-Table S7**). Remarkably, although co-optimizing binding affinities alongside concentrations provides greater parameter flexibility per monomer, our natural bZIP clusters, despite being strictly constrained by fixed, evolutionarily determined *K_d_* values, still achieved near-maximal computational versatility (15 out of 16 gates) using compact natural communities.

This drastic reduction in required molecular components (compared with random-affinity frameworks) implies that natural protein interactomes possess highly dense architectures sculpted by evolutionary pressures^18^. Rather than presenting arbitrary binding profiles, biological families harbor a vast, latent computational capacity. This intrinsic complexity allows natural networks to support sophisticated signal processing out-of-the-box, offering a compact, scalable design paradigm for synthetic biology that circumvents the massive bottlenecks of *de novo* protein design^16,17^.

Beyond overall network scale, a fundamental question concerns the underlying biochemical drivers that enable certain individual networks to compute complex, non-linearly separable operations while others fail. We hypothesize that several candidate mechanisms shape these divergent computational behaviors, including competitive binding, where multiple monomer species compete for a shared dimerization partner; molecular sequestration, in which a monomer is actively trapped within a high-affinity dimer and rendered unavailable for alternative interactions; species redistribution, driven by input-induced shifts that dynamically reorganize the global monomer pool that contributes to the formation of the output dimer; and thermodynamic dominance, where local balances between *K_d_* values and baseline concentrations dictate which dimeric species predominate at equilibrium.

On the other hand, analyzing which specific logic gates are missing or most challenging across our network population offers critical insight into the physical limits of mass-action biocomputing.

Across all evaluated individual networks, XNOR emerged as the least frequently computed function, followed closely by XOR (**Supplementary note S2-Table S5**). This difficulty directly reflects their non-linearly separable nature, which demands a highly delicate balance of competitive sequestration and monomer displacement across multiple truth-table states. Crucially, neither gate was completely absent from our dataset: at least one individual network successfully implemented XNOR or XOR. This demonstrates that while non-linear operations represent the primary computational bottleneck for competitive dimerization, they do not constitute an absolute thermodynamic impossibility. Instead, their execution relies on highly specific topological features and *K_d_* distributions, though in certain cases, missing solutions may also reflect the boundaries of the heuristic search where the genetic algorithm failed to converge on an existing, narrow solution space.

These results establish that natural bZIP interaction networks support a broad repertoire of two-input logic, including XOR and XNOR. Achieving up to 15 of the 16 Boolean functions shows that experimentally measured interactions provide sufficient flexibility for diverse computations through accessory concentration tuning and output-dimer selection. This directly addresses the affinity-design bottleneck by replacing abstract interaction requirements with identifiable proteins and defined configurations.

### 2.3. Impact of Input Monomer Selection on Logical Versatility

After evaluating the computational versatility of each natural bZIP cluster, we next examined whether the selection of input monomers influenced the number of logic gates that a cluster could compute. In the initial optimization experiments, input monomers were selected based on their connectivity within the cluster (**Section 5.6.1**). This choice was motivated by the assumption that highly connected monomers would provide greater computational capacity, since input perturbations introduced through these nodes could be distributed through a larger number of competitive interactions.

We performed a complete input-pair analysis for *N. vectensis* cluster #0 at 4°C, where we optimized the accessory monomer concentrations for all 21 possible pairwise input combinations within the cluster and evaluated the maximum number of target logic gates achieved by each configuration. The results of this exhaustive screening are summarized in **Supplementary note S4-Table S8** and visualized in **Figure 4**.

**Fig. 4.**
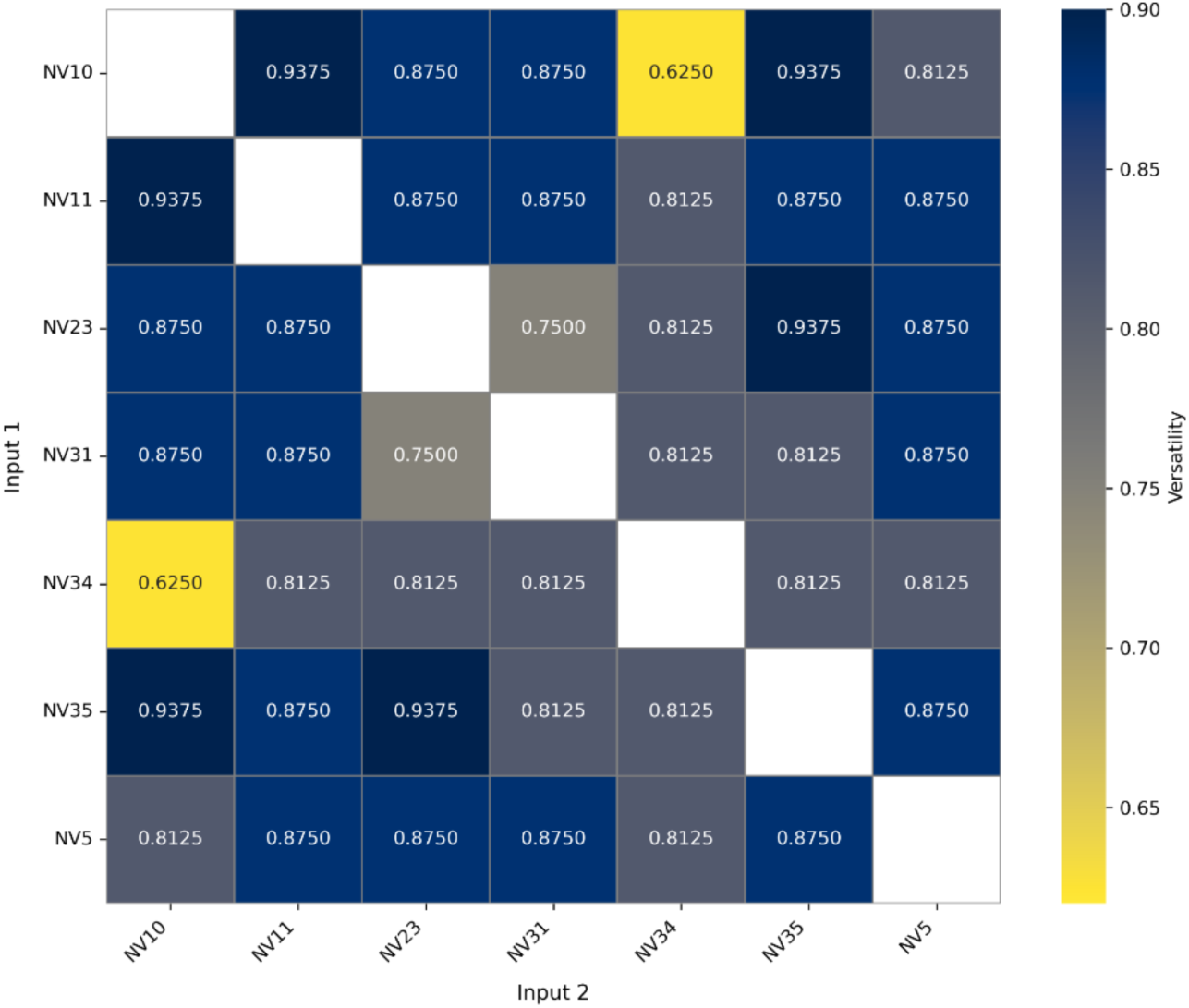
Computational versatility across all pairwise input combinations in *Nematostella vectensis* cluster #0 (4 °C). Heatmap showing the fraction of 16 target two-input logic gates successfully computed by each pairwise input monomer combination. Values range from 0.6250 (10/16 gates, yellow) to 0.9375 (15/16 gates, dark blue).

For each input pair, the total input connectivity was calculated as the combined number of connections from both input monomers, corrected by the subtraction of repeated interactions (**Table S8**). The resulting versatility values show that the computational output of the cluster depends strongly on which monomers are selected as inputs, but not in a simple monotonic way with respect to their total number of connections (**Figure 4**).

The empirical data demonstrates that logical versatility fluctuates significantly depending on the specific identity of the selected inputs rather than their raw connectivity metrics. For instance, pairing two highly connected monomers such as **NV31** (7 connections) and **NV23** (7 connections) yields a cumulative input edge footprint of 13 unique links, yet its functional performance is restricted to a relatively low versatility of 12 out of 16 gates (0.75). Conversely, pairing a weakly connected monomer like **NV11** (3 connections) with an intermediate node like **NV10** (5 connections) results in a total input footprint of only 8 unique links, yet it achieves a near-maximum versatility of 15 out of 16 gates (0.9375) (**Table S8**). These results indicate that input connectivity alone does not determine the computational versatility of the cluster.

The heatmap in **Figure 4** highlights this input-dependent behavior. Rather than showing a uniform increase in versatility with input degree connectivity, the matrix reveals that specific monomers tend to produce higher or lower computational performance depending on their pairing. In particular, **NV10** and **NV35** appeared frequently in highly versatile input combinations, whereas **NV34** was associated with lower versatility across several pairings. This suggests that some monomers are more effective as input channels, while others may constrain the ability of the cluster to generate separable output patterns.

#### 2.3.1 Discussions on the Impact of Input Monomer Selection

These findings prove that raw physical connectivity (node degree) is not the primary determinant of computational latency within natural protein interaction networks. If the absolute number of interactions does not dictate logical versatility, other subtle structural and thermodynamic properties of the graph topology must govern the distribution of chemical equilibrium states.

We hypothesize that this behavior may be driven by higher-order topological metrics, such as betweenness centrality, which quantifies how often a node acts as a bridge along the shortest paths between other members of the community, or the presence of localized topological bottlenecks^31,32^. Furthermore, the explicit distribution of the individual equilibrium dissociation constants (*K_d_*) linked to these specific monomers likely creates precise thermodynamic regimes that can either amplify or quench the competitive binding landscape. A complete understanding of which network descriptors drive these computational enhancements remains a critical objective for future structural analysis and experimental validation.

These results establish input selection as a consequential design variable alongside accessory concentration tuning and output-dimer selection. The comparison provides a practical basis for prioritizing input pairs and avoiding configurations that underuse the available interaction network. By identifying more versatile assignments within an existing protein assembly, this analysis guides the design of molecular processors that exploit natural interactions more effectively before resorting to additional components or protein engineering.

### 2.4. Three-Input Logic Computation

A critical challenge in the development of molecular diagnostic platforms and intelligent cell therapies is the need to process higher-order biological signal combinations without increasing system complexity. While two-input operations demonstrate basic decision-making, real-world cellular environments demand the integration of multiple distinct disease biomarkers simultaneously^33,34^. Scaling synthetic genetic biocomputing traditionally requires constructing multi-layered cascades, which increases metabolic burden^35^, slows system response, and amplifies noise under certain conditions^36^. An assessment of whether unengineered, fixed-affinity protein interactomes can scale toward three-input logic within a single competitive layer addresses this operational challenge directly, bypassing both *de novo* protein design and the architectural burden of multi-step genetic or enzymatic cascades.

After demonstrating that natural bZIP clusters can compute a broad range of two-input Boolean functions, we next evaluated whether the most versatile clusters could scale toward more complex three-input computation. For this analysis, we selected the clusters that showed high computational performance in the previous two-input experiments (*N. vectensis* cluster #0, *N. vectensis* cluster #3, and *C. intestinalis* cluster #7) and subjected them to continuous genetic algorithm optimization across a targeted benchmark suite of 17 three-input logic functions (**Section 5.6.2**). Because the input-selection analysis showed that the identity of the input monomers can influence cluster versatility, we tested different combinations of input monomers within each selected cluster. To reduce computational cost, we restricted this analysis to three input connectivity categories: highly connected, intermediately connected, and poorly connected monomers. From these trials, we retained the individual networks with the highest three-input versatility.

The empirical optimization benchmarks (**Table 3**) demonstrate that natural protein interaction networks are capable of scaling to three-input logic functions without requiring structural or synthetic modifications. Consistent with its high node density and extensive internal cross-talk, *N. vectensis* cluster #3 emerged as the most versatile architecture, successfully evaluating 13 out of the 17 target logic gates (76.47% logical versatility). It was followed by the 9-monomer *C. intestinalis* cluster #7 (11/17 gates, 64.7%) and the 7-monomer *N. vectensis* cluster #0 (9/17 gates, 52.9%). **Supplementary note S2-Table S6** summarizes which of the 17 three-input logic functions in the proposed benchmark suite were successfully computed by each cluster.

**Table 3.** Computational performance and structural metrics of natural bZIP clusters evaluated across the 17-function 3-input logic benchmark suite. For each cluster, the monomer count, active interaction footprint, number of resolved logic functions, total evaluated operations, and resulting versatility score are reported.

| Organism | Cluster | Number of monomers | Number of possible dimers / interactions | Number of active dimers / interactions | Connection density | Computed logic gates | Theoretical two input logic gates | Versatility |
|---|---|---|---|---|---|---|---|---|
| **NV** | #0 | 7 | 28 | 20 | 0.71 | 9 | 17 | 0.5294 |
| **NV** | #3 | 13 | 91 | 41 | 0.45 | 13 | 17 | 0.7647 |
| **CI** | #7 | 9 | 45 | 29 | 0.64 | 11 | 17 | 0.6470 |

Our results indicate that clusters with high two-input versatility can retain substantial computational capacity when extended to three-input logic, although the transition to higher-order input spaces reduces the number of functions that can be successfully implemented.

Most of the linearly separable or threshold-like gates were computed by the three selected clusters. This suggests that the optimized dimerization networks can still reproduce input-output behaviors that depend on broad activation patterns across three inputs. However, the more complex non-linearly separable functions were substantially harder to compute. Among these functions, only EXACTLY ONE was computed by the most versatile cluster. In contrast, the remaining non-linearly separable or highly combinatorial functions were not consistently achieved by the selected networks.

#### 2.4.1 Discussions on Three-input logic gates

Scaling molecular computation from two to three inputs represents a major transition in logical complexity^17^ as the underlying chemical system must discriminate between eight distinct input combinations rather than four. While these natural networks successfully executed a significant fraction of the target operations, up to 76.5% versatility in the largest cluster, their computational performance exhibited a clear boundary: they favored linearly separable and threshold operations but largely failed to resolve highly non-linear functions.

This distinct performance boundary reveals the physical constraints that shape unengineered, natural biochemical computation. As the number of independent input variables increases from two to three, the combinatorial space of equilibrium chemical states expands exponentially. To resolve a highly non-linear three-input gate (such as XOR or XNOR) at chemical equilibrium, the underlying interaction topology must dynamically redistribute free monomer populations across eight distinct truth-table quadrants. In our empirical trials, three-input computational versatility scaled directly with cluster size and internal connectivity. However, across all three substrates, successful execution was predominantly restricted to linearly separable and threshold-like operations, with only a single non-linearly separable function (EXACTLY ONE) successfully resolved by the largest cluster (*N. vectensis* #3).

To contextualize these natural limits, it is insightful to compare our findings with synthetic biocomputing frameworks where binding affinities are not constrained by natural evolution. Parres-Gold *et al.*^9^ evaluated three-input logic in synthetic networks of 4 to 10 monomers by co-optimizing both the pairwise association constants (*K_a_*) and monomer concentrations. When mapped onto our 17-function benchmark suite, their framework successfully resolved 9 logic gates. Crucially, because Parres-Gold *et al.*^9^ allowed *K_a_* values to vary freely across an unconstrained parameter space, their model was capable of computing complex, non-linearly separable three-input operations (such as three-input XOR and XNOR). The remaining functions in our suite, specifically the six single-variable Buffers and Inverters (NOT A, NOT B, NOT C, Buffer A, Buffer B, and Buffer C), were not identified in their study simply because they were excluded from their targeted search, which focused exclusively on fundamental multi-input Boolean operations. Similarly, in our prior work^14^ on minimal 4-monomer synthetic networks where both *K_a_* and concentrations were co-optimized, we identified solutions for 10 out of the 17 benchmark gates in the present suite; we notably discovered configurations capable of implementing the highly non-linear three-input XNOR gate.

A broader methodological challenge in comparing molecular logic across different studies is the current lack of a universal consensus regarding which specific functions should comprise a standard three-input benchmark suite, beyond the general agreement that computing non-linearly separable gates represents the ultimate benchmark of computational capacity^17,37^. Nevertheless, juxtaposing these co-optimized synthetic models against our natural bZIP networks highlights a fundamental design trade-off in biomolecular computing. When binding affinities can be freely engineered *de novo*, small synthetic networks (4–10 monomers) can readily access highly complex non-linear state spaces. In contrast, when operating with fixed, natural *K_a_* values, competitive concentration tuning alone allows natural bZIP families to achieve substantial computational capacity for linear and threshold logic (up to 13/17 gates) using compact, non-engineered clusters. However, transitioning these unengineered natural networks toward advanced non-linear three-input operations will probably require implementing multi-layered molecular cascades or applying targeted synthetic engineering to sculpt the underlying binding affinity landscape.

In summary, the successful computation of up to 76.5% of the three-input benchmark suite demonstrates that unengineered protein interactomes can effectively process multi-signal molecular environments without the need for complex, multi-layer genetic architectures. By showing that natural concentration tuning alone is sufficient to integrate three distinct inputs into linear and threshold decisions, these findings prove that biological networks provide a compact substrate for real-time diagnostic evaluation. While highly non-linear three-input operations appear to mark the physical boundary of single-layer fixed-*K_d_* networks, achieving robust multi-input processing in non-engineered clusters successfully bridges the gap between theoretical biocomputing and practical applications.

### 2.5. Sensitivity Analysis and Tolerance to Expression Noise

A fundamental hurdle in translating synthetic biocomputing systems to biological environments is their vulnerability to intrinsic molecular fluctuations^38^. While deterministic model predictions validate theoretical logic execution under nominal conditions, real-world cellular and cell-free environments are inherently noisy, characterized by transcriptional bursting, expression variability, and fluidic delivery noise^39^. A logic circuit that computes flawlessly under static concentrations is practically useless if minor molecular variations degrade its decision boundaries. An evaluation of the functional resilience of our unengineered natural bZIP networks under stochastic perturbations addresses this problem directly. Evidence that fixed-affinity competitive interactomes maintain distinct, non-overlapping logic states under realistic noise levels proves that natural protein families harbor intrinsic thermodynamic buffering and confirms their viability as robust, fault-tolerant molecular processors.

To verify that our two-input 135 individual networks maintain predictable Boolean computation in fluctuating biological environments, we subjected each circuit to systematic concentration perturbations. To capture realistic gene-expression noise and transcriptional bursting, all monomer abundances were perturbed using a gamma distribution noise protocol across six progressive intensities (CV = 0.05 to 0.50) (**Section 5.7.1**). To decouple simple nominal performance from true biochemical resilience, network behavior was quantified applying three complementary metrics (**Section 5.7.2**): Probability of error (*P_error_*), measures the frequency of logical failure under concentration perturbations; signal separation (S) quantifies the separation between logical-zero and logical-one output distributions; and logarithmic root-mean-square deviation (RMSD) measures how far the perturbed output profile deviates from its nominal response on a logarithmic scale. Together, these metrics distinguish the preservation of logical function from changes in output concentrations.

#### 2.5.1. Noise Sensitivity and State Separation Boundaries

The probability of error (*P_error_*) revealed marked heterogeneity in noise tolerance across the perturbed network population (**Fig. 5A**). The error profiles did not increase uniformly but instead distinguished two clear functional classes: robust networks, which remained virtually error-free even at CV = 0.50, and fragile networks, which computed the target gate under nominal conditions but suffered progressive logical failure under minor concentration fluctuations. This disparity underscores that nominal performance alone is insufficient to validate a biological logic module^40^.

**Fig. 5.**
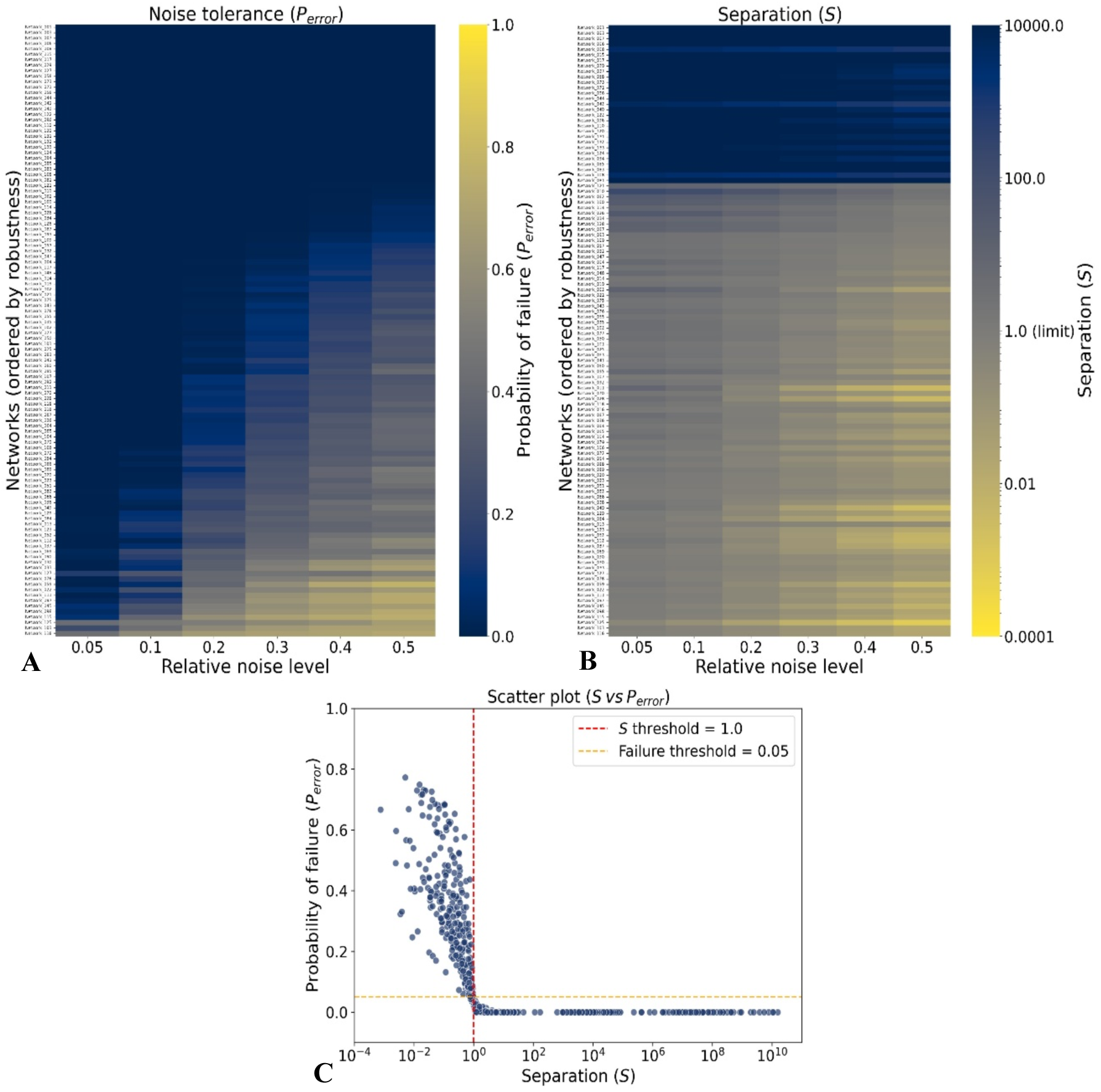
Robustness of optimized bZIP individual networks under stochastic concentration noise. A) Heatmap of probability of failure (*P_error_*) across increasing relative noise levels. Each row represents an individual network, and color intensity indicates the probability of logical failure. B) Heatmap of signal separation (S) across the same noise levels, where higher values indicate stronger separation between logical-zero and logical-one output states. In both heatmaps, darker blue regions denote more robust network behavior. C) Scatter plot of *P_error_* as a function of S, with reference thresholds at S = 1.0 and *P_error_* = 0.05 defining the separation boundary and target error tolerance used to identify robust logic behavior.

To explain why some networks failed while others remained stable, we analyzed the separation between logical-zero and logical-one output states (**Fig. 5B**). Robust networks consistently preserved a clear, non-overlapping signal window between binary classes. The correlation of S against *P_error_* uncovers a highly non-linear phase transition in system performance (**Fig. 5C**). This correlation confirms that robust logic operations rely heavily on the magnitude of the signal window between binary states. When the separation window is large, stochastic concentration bursts cannot easily bridge the gap between states and the integrity of the output is preserved. When the margin between states is narrow, even small concentration variations can blur the boundary and compromise the logic gate representation. Individual networks that exhibit a wide signal window (S > 1.0) concentrate tightly near a zero-error rate. However, as soon as the signal distributions overlap (S < 1.0), signal resolution is lost, the data points scatter broadly, and failure rates increase abruptly. This mathematical boundary demonstrates that robust logic operations rely strictly on maintaining a wide separation window to prevent stochastic concentration bursts from bridging the gap between binary states, preserving the integrity of the output ^38,41^.

#### 2.5.2. Network Population Dynamics and Output Stability

While *P_error_* and S capture binary success thresholds, they do not measure the continuous structural deformation of the output profile relative to its optimized nominal baseline. To track this trajectory, we monitored the continuous *RMSD_lo_*_g_ deviation^42^ (**Section 5.7.2**).

Representative networks are compared in **Figure 6A**, including the three most robust networks (relative to *P_error_*), Network_001, Network_003, and Network_007, and the least robust network, Network_116. The x-axis represents the relative noise level, and the y-axis represents *RMSD_lo_*_g_. The lines indicate the central tendency, while the shaded regions represent the interquartile range. As expected, *RMSD_lo_*_g_ increased progressively with the noise level, which indicates that output profiles diverge from their nominal values as perturbations become stronger. However, the magnitude and dispersion of this deviation depended strongly on the network. The most robust network (Top 1) exhibited low *RMSD_lo_*_g_ values and a narrow interquartile range, indicating that its output remained close to the nominal response under perturbation. In contrast, the second most robust network (Top 2) showed larger deviations and broader interquartile ranges. Interestingly, even the least robust network exhibited lower *RMSD_lo_*_g_ values and narrower interquartile ranges than Top 2, implying that smaller deviations from the nominal response did not necessarily correspond to greater robustness. Crucially, this analysis highlights why the evaluation of all three statistical metrics simultaneously is necessary. An individual network can experience a high continuous deviation RMSD, yet successfully avoid a logical failure (*P_error_* = 0) as long as the underlying competitive binding architecture keeps the output distributions from overlapping (S > 1.0).

**Fig. 6.**
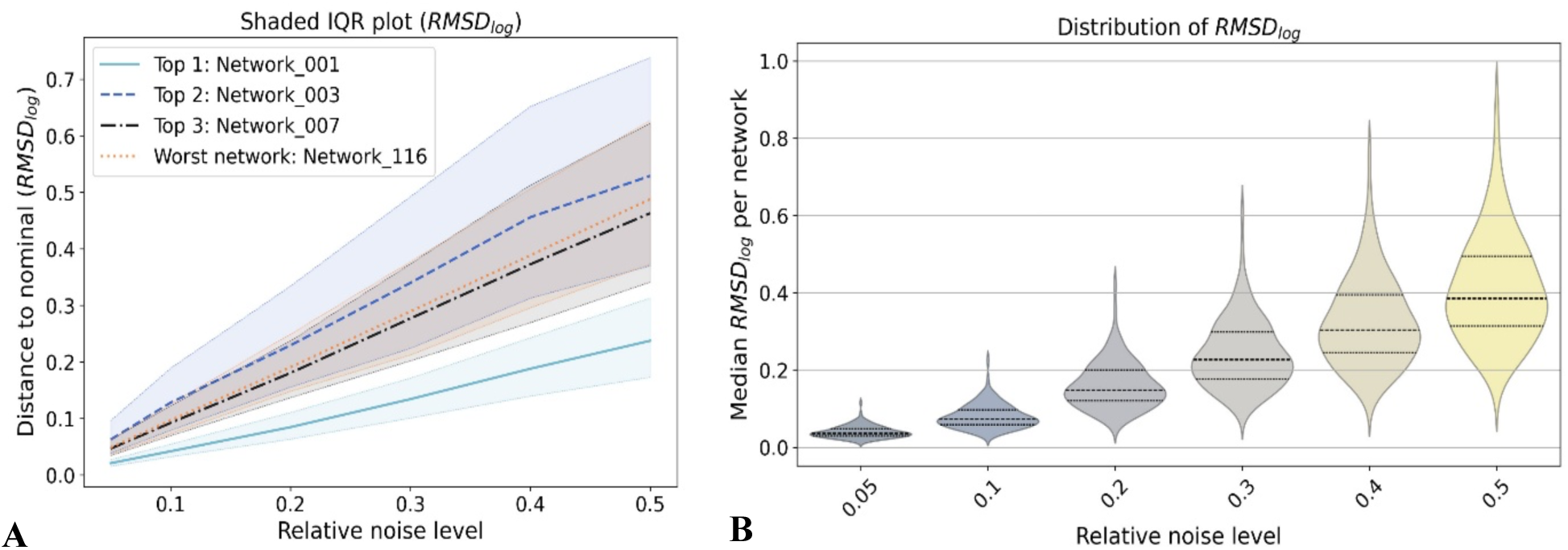
Output-profile stability of optimized bZIP individual networks quantified by *RMSD_lo_*_g_. A) Representative *RMSD_lo_*_g_ trajectories across increasing relative noise levels. Lines denote median deviations from the nominal optimized output profile, and shaded regions indicate interquartile ranges across perturbation trials. B) Violin plots summarize the population-wide distribution of *RMSD_lo_*_g_ across all optimized individual networks at each noise level, showing how output-profile dispersion increases under stronger perturbations.

To observe how these structural variations evolve across the entire population of simulated networks, we examined the full distribution of continuous errors (**Fig. 6B**). The violin plots indicate that the median *RMSD_lo_*_g_ increases with relative noise level, and that the dispersion of responses also becomes broader as perturbation increases. At high noise levels, longer upper tails indicates that some networks are substantially more sensitive than others. This suggests that, under low noise, most networks behave relatively similarly, whereas at high noise levels the differences between robust and sensitive networks become more pronounced^41^.

#### 2.5.3. Discussions on computational robustness

These sensitivity analyses provide key design rules for synthetic biology and molecular diagnostics^42^. Although the division of the networks in two classes (robust and fragile) can be partially explained by the existence of two fitness thresholds (**Section 5.6.1**), there are cases where an individual network with high fitness has also a high probability of error. For example, for network #059 with a fitness of 0.80. At the lowest noise level, the network maintains a very low probability of failure (0.06) but as soon as the noise increases the probability of error drastically grows reaching 0.773 for the highest noise level. This individual network would therefore be classified as fragile despite its high fitness score, which suggests that network robustness depends on factors beyond fitness alone. High-robustness configurations may correspond to specific bZIP dimerization topologies where internal competitive binding and distinct affinity preferences form a biochemical buffer that absorbs fluctuations in individual monomers^38,43^. For practical *in vivo* or *in vitro* applications where absolute molecular concentrations cannot be perfectly controlled, networks maintaining *P_e_*_rror_ < 0.05 and S > 1.0 represent viable candidates for reliable biocomputing^40,44^.

Importantly, this robustness must be interpreted strictly as an intrinsic property of the thermodynamic models under simulated conditions, rather than evidence that these bZIP protein families naturally function as digital Boolean gates within their native cellular environments. Instead, these outcomes confirm that natural protein-protein interaction networks possess a powerful, latent computational robustness that can be strategically harnessed via optimized concentration frameworks to build reliable synthetic circuits^30^.

Ultimately, the identification of a robust subset of unengineered networks that preserve tight decision boundaries (*P_error_* < 0.05, S ≫ 1.0) under severe noise solves the critical problem of biological variability and expression fluctuations. These results establish that *de novo* protein engineering is not required to achieve fault-tolerant molecular logic. This noise-filtering capability provides a quantitative baseline for the selection of highly resilient candidate circuits and supports the development of predictable, failure-resistant processors that can function accurately in fluctuating *in vivo* and *in vitro* environments.

## 3. Biological Constraints and Thermodynamic Scaling Frameworks

The transition from theoretical modeling to real-world deployment requires addressing the concrete physical limits of biological systems^45^. While our simulations validate that unengineered bZIP interactomes can evaluate complex Boolean logic at physiological nanomolar scales, translating these molecular circuits into fast, cell-free experimental assays introduces operational challenges, such as signal detection limits^46^, fluidic handling constraints^47^, and fixed biological affinity bounds^18^. A theoretical circuit design is only as useful as its capacity to be physically realized in experimental platforms. The definition of these biological boundaries and the formulation of a robust thermodynamic scaling framework directly address this translational issue. Establishing how nanomolar predictions map onto micromolar or millimolar experimental regimes guarantees that our fixed-affinity, concentration-tuned networks can maintain logical accuracy and signal contrast during physical *in vitro* implementation and provides a clear engineering roadmap for real-time diagnostic applications.

### 3.1. Experimental Resolution Bounds of Natural Binding Parameters

The first limitation arises from the bounded parameter space of the empirical dataset. In our simulations, association constants (*K_a_*) were constrained to the experimental resolution range reported by Reinke *et al.*^18^, spanning from 2 × 10^−4^ *nM*^−1^ (*K_d_* = 5000*nM*) to 1*nM*^−1^ (*K_d_* = 1*nM*). While the lower bound (*K_a_* ≈ 10^−3^ *nM*^−1^) does not impose a critical constraint, since such weak affinities yield virtually negligible dimerization under equilibrium mass action, the upper bound (*K_d_* = 1*nM*) represents a physical measurement ceiling of the experimental assay rather than an absolute biological limit. In contrast, theoretical models, such as those by Parres-Gold *et al.*^9^, considered synthetic association constants up to 10^7^ *nM*^−1^. Consequently, the empirical dataset leaves a blind spot regarding potentially ultra-high-affinity natural interactions (*K_d_* < 1*nM*) that may exist in living cells but remain unquantified.

### 3.2. Deviation from Ideal Dimerization Regimes

The fixed nature of natural binding affinities also causes our system to operate outside the ideal thermodynamic dimerization boundaries established in unconstrained synthetic studies. In theoretical frameworks^9,14^, the interaction affinity range (*K_a_* ∈ [10^−5^, 10^7^] *nM*^−1^) and concentration interval (C ∈ [10^−3^, 10^3^]*nM*) are chosen to guarantee two ideal asymptotic states:

1. The strongest affinity at the lowest concentration yields 99% dimerization.
2. The weakest affinity at the highest concentration yields only 1% dimerization (negligible background).

In our model, input and accessory concentrations were restricted to physiologically realistic cellular expression levels (10^−3^*nM* to 10^2^*nM*)^48^. This range corresponds to abundances from fewer than five to tens of thousands of protein copies per cell^49^. Because natural bZIP *K_a_* values are fixed and capped at 1*nM*^−1^, the system departs from these ideal theoretical limits:

- Strongest affinity (*K_a_* = 1*nM*^−1^) at lowest concentration (10^−3^*nM*): Achieves only 0.098% dimerization, much lower than the ideal 99% threshold.
- Strongest affinity (*K_a_* = 1*nM*^−1^) at highest concentration (10^2^*nM*): Yields 90% dimerization, slightly below the ideal asymptotic ceiling.
- Weakest affinity (*K_a_* = 2 × 10^−4^*nM*^−1^) at highest concentration (10^2^*nM*): Yields 2% dimerization, closely matching the ideal 1% background target.
- Weakest affinity (*K_a_* = 2 × 10^−4^*nM*^−1^) at lowest concentration (10^−3^*nM*): Yields 0% dimerization, an ideal baseline behavior.

Importantly, despite operating well outside the ideal theoretical dimerization regime due to these fixed biological parameters, our optimization framework proves that competitive equilibrium networks successfully preserve robust Boolean logic discrimination.

### 3.3. Concentration Scaling Framework for Experimental *In Vitro* Implementation

A practical consideration for experimental translation is the concentration regime. While our simulations operate in the nanomolar range (10^−3^*nM* to 10^2^*nM*) to mirror physiological intracellular abundance, cell-free or *in vitro* diagnostic assays often require higher concentration scales (micromolar *μM* or millimolar *mM*) for optical detection, fluidic handling, and signal-to-noise optimization^50^.

To map nanomolar predictions onto higher experimental scales without altering logic gate performance, we formulate a non-dimensional mass-action framework based on chemical reaction kinetics and dimensional analysis^51,52^. Chemical equilibrium under mass action [*D_ij_*]_*eq*_ = *K_ij_*[*M_i_*]*_eq_* [*M_j_*]*_eq_* can be non-dimensionalized using a characteristic scale 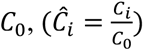, which shows that species distributions depend strictly on the dimensionless affinity product 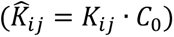,. Scaling all total loaded species abundances uniformly by a factor *α* (*C_tot_*′ = *αC_tot_*) yields two operational strategies:

**Scenario A (Exact Fractional Invariance):** Adjusting environmental buffer conditions or temperature to scale physical affinities inversely 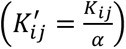 keeps 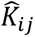, invariant. This guarantees that equilibrium fractional species distributions ([*D_ij_*]*_eq_* / *C_total_*) and decision boundaries remain 100% identical to simulation predictions.

**Scenario B (Fixed-Affinity Quadratic and Linear Scaling):** When binding affinities remain fixed 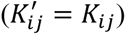,, scaling species abundances upward by *α* increases the dimensionless product. In low-conversion or basal output background regimes (*x_i_* ≈ 1), dimeric outputs scale quadratically ([*D_ij_*]*_eq_*′ ≈ *α*^2^[*D_ij_*]*_eq_*) across input and accessory pairings. Conversely, in highly saturated regimes producing high output signal levels (*x_i_* → 0), dimer output scaling transitions smoothly from quadratic (*α*^2^) to near-linear (≈ *α*).

This regime transition produces a favorable sigmoidal compression effect: it suppresses low-concentration basal output background noise and maintains high output signal levels, which expands the absolute signal margin (Δ[*D_output_*]) for cell-free detection. Complete mathematical proofs, non-dimensionalization derivations, and parameter-space validity analyses are detailed in **Supplementary Note S5**.

By establishing this non-dimensional scaling framework, we effectively narrow the gap between nanomolar theoretical modeling and physical, micromolar-scale *in vitro* deployment. These mathematical formulations prove that the empirical limits of the binding dataset and departures from ideal dimerization regimes do not invalidate logical computation. Instead, concentration-tuned natural bZIP networks preserve their decision boundaries under environmental buffer tuning (Scenario A) or even benefit from an expanded signal-to-noise margin (Δ[*D_output_*]) via non-linear regime transitions under fixed affinities (Scenario B). The resolution of this translational bottleneck guarantees that our non-engineered biocomputing designs can be directly ported into real-time, cell-free experimental assay platforms without compromising logical fidelity.

## 4. Conclusions and Future Work

In this study, we established a systematic optimization and perturbation framework to map the latent computational capacities of natural, unengineered protein dimerization networks. By evaluating the thermodynamic equilibrium of basic bZIP communities from *Nematostella vectensis*, *Ciona intestinalis*, and *Homo sapiens*, we demonstrated that complex multi-input processing can emerge directly from single-layer biological systems without structural modifications or slow transcription-translation dynamics. This work provides a critical proof-of-concept that evolutionarily tuned interactomes harbor built-in computational power, offering a ready-to-use alternative to the daunting bottlenecks of *de novo* protein engineering^16,17^.

Within the scope of our simulations, the curated natural bZIP clusters exhibited a remarkably favorable logical density. While previous frameworks based on randomly assigned binding profiles required a minimum of 20 monomers to resolve 10 multi-input functions^9^, our selected natural subsets successfully executed up to 15 out of 16 distinct two-input logic gates, and up to 13 functions across a 17-gate three-input benchmark suite, using constrained networks of only 9 and 13 monomers. This performance confirms that the evolutionary topology of natural protein families inherently supports rich competitive binding regimes that efficiently process combinatorial signal inputs.

Harnessing this structural efficiency directly addresses pressing obstacles in synthetic biology: the slow response times of genetic circuits and the single-input limitations of conventional biosensors^8^. The natural networks evaluated here overcome these constraints by processing multi-signal signatures within seconds. Furthermore, stochastic perturbation analysis under Gamma-distributed noise confirmed that this computational capacity is resilient to gene-expression fluctuations. A significant subset of optimized networks absorbed severe concentration variability (CV = 0.50) while maintaining wide signal separation margins (S ≫ 1), providing a fault-tolerant foundation for real-world biological deployments. Finally, our non-dimensional thermodynamic scaling framework bridges the operational gap between nanomolar simulation scales and micromolar *in vitro* cell-free assays, proving that decision boundaries remain intact or even benefit from noise suppression during physical implementation.

It is important to emphasize that model-predicted concentrations and candidate physical proteins serve as a theoretical baseline rather than empirical validation. However, this framework successfully generates robust, quantitatively defined circuit architectures ready for experimental evaluation.

For these reasons, the immediate next step for this research is to perform rigorous *in vitro* experimental assays using recombinant bZIP proteins from the selected clusters. Guided by our non-dimensional scaling principles, future studies will evaluate whether cell-free reactions replicate predicted steady-state output dimer profiles under physical titration conditions. Additionally, exploring how these networks behave within interconnected molecular cascades, or mapping their topologies to higher-order graph-theoretic descriptors, will deliver essential design rules to harness natural protein architectures as robust, real-time biocomputing processors.

## 5. Methodology

### 5.1. Theoretical Framework of Dimerization Networks and Chemical Equilibrium

In a dimerization network, the reversible association of two monomers into a dimer acts as an input-output biochemical computation^55^. Under this framework, molecular species are classified into three distinct computational roles: input monomers, whose initial concentrations represent the input signals; accessory monomers, which provide the competitive background required to shape the network’s behavior; and output dimers, which are the specific dimers whose equilibrium concentrations are tracked as the computational output. Because any given network forms multiple unique dimers at equilibrium, the concentration of any specific dimer can be tracked as the output signal.

This system is governed by the law of mass action. At chemical equilibrium, the dynamic association and dissociation of monomers *M_i_* and M_j_ into the dimer *D_ij_* (Eq. 1) is quantified by the association equilibrium constant, *K_ij_* (Eq. 2):

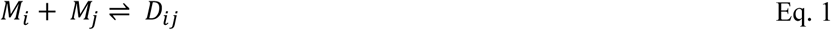

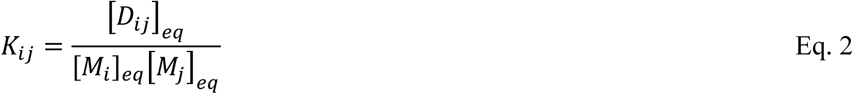

Here, [*D_ij_*]*_eq_*, [*M_i_*]*_eq_*, and [*M_j_*]*_eq_* denote the respective equilibrium concentrations of the dimer and the free, unbound monomers.

To simulate these networks, we implemented a thermodynamic framework using the Python library, Equilibrium Toolkit (EQTK)^28,29^. By solving the governing equations derived from reaction stoichiometry, equilibrium constants, and mass conservation, EQTK determines the unique steady-state concentrations of all species. This allows us to compute the concentration of any target output dimer under specific input conditions and characterize the network’s logical behavior.

### 5.2. Data Curation and Extraction: The Reinke Dataset

To ground our computational analysis in experimentally measured interaction parameters, we utilized the dataset reported by Reinke *et al.*^18^, which comprehensively quantifies the equilibrium dissociation constants for basic leucine zipper (bZIP) interaction networks across multiple organisms. This naturally occurring protein family provides a robust, experimentally bounded parameter space for testing biological computation.

The dataset reports *K_d_* values for bZIP families across multiple organisms measured at 4°C, 21°C, and 37°C. The reported dissociation constants span an experimental range from 1 to 5000 *nM*, where values beyond these thresholds reflect the resolution limits of the binding assay rather than true physical bounds.

For compatibility with EQTK, the reported dissociation constants were converted into association constants, *K_a_*, via **Eq. 3**:

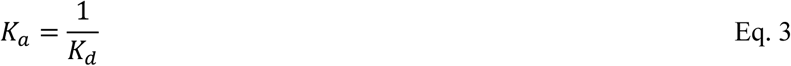

After this transformation, the resulting *K_a_* values ranged from 2 × 10^−4^*nM*^−1^ to 1*nM*^−1^.

Integrating this dataset into EQTK required a symmetric interaction matrix, where the binding affinity of monomer *i* to monomer *j* (*K_ij_*) equals that of monomer *j* to monomer *i* (K_*ji*_). Because Reinke *et al.*^18^ measured interactions pairwise in separate experimental runs, their raw data yielded asymmetric matrices (*K_ij_* ≠ K_*ji*_). To resolve these discrepancies, we cross-referenced the experimental correlation coefficients reported for each measurement and selected the affinity value (*K_ij_* or K_*ji*_) with the higher correlation coefficient. This procedure yielded a fully symmetric interaction matrix for each bZIP network, which was subsequently used as input for EQTK to compute the equilibrium concentrations of all molecular species.

### 5.3. Topological Network Analysis and Cluster Extraction

To construct a natural computational substrate with a high density of interactions, we focused on the bZIP interaction networks of three specific organisms from the Reinke *et al.*^18^ dataset: *Homo sapiens*, *Nematostella vectensis*, and *Ciona intestinalis*. These organisms were chosen because they exhibit the most highly connected networks within the empirical dataset. To capture weak but computationally valuable interactions, we primarily focused on datasets measured at 4°C. Additionally, a human bZIP cluster at 37°C was included to assess performance under physiological temperatures.

While this comprehensive topological profiling proved that modeling full networks was theoretically possible, we ultimately discarded this approach for practical applications. Since one of the long-term goals of this work is experimental implementation, using complete networks would impose a substantial metabolic burden in living systems^23^ and would also introduce excessive complexity in cell-free platforms^56^. For this reason, we focused on extracting smaller natural subnetworks, here referred to as clusters.

We first evaluated whether the networks were organized around highly connected hubs. Consistent with the observations reported by Reinke and Amoutzias^12,18^, the bZIP networks showed scale-free organization, which indicates that clusters could not be defined solely on the basis of hub-centered structures. Therefore, clusters were identified using the Louvain community detection algorithm in the python-louvain library. The Louvain method^25^ is a fast heuristic approach that identifies clusters by maximizing network modularity, a metric that compares the density of internal connections within a group against expected random connections to the outside. The optimization was weighted by the experimental *K_a_* values, with *random state* = 42 to ensure reproducibility and *resolution* = 1 to retain the standard modularity formulation. NetworkX^57^ in python was used to construct the graphs corresponding to each cluster.

After community detection, each cluster was analyzed individually. For the computational analysis, clusters were selected to represent a broad range of network sizes. Small clusters of three monomers represented the minimum configuration required to accommodate two input monomers and one accessory monomer. Intermediate clusters of six to nine monomers offered a balance between interaction complexity and the computational cost of genetic algorithm optimization. A large cluster of 13 monomers was included to assess whether greater network size was associated with increased computational capacity relative to intermediate-sized clusters. To cover this size range while keeping computational costs manageable, 13 representative clusters across the three organisms were selected from the eight clusters identified per species.

For each selected cluster, we quantified monomer interactions along with the mean and standard deviation of their binding strengths. Because Louvain partitioning extracts densely integrated subgraphs by maximizing internal modularity, the extracted bZIP clusters form strongly connected topological modules, thereby facilitating effective signal propagation between inputs and output dimers.

The analysis distinguishes three levels of organization. At the organism level, all bZIP proteins and interactions included in the dataset constitute the complete network. Subsets extracted from this network are termed clusters. When a cluster is assigned specific input monomers, accessory concentrations, and an output dimer that together enable at least one logic function, the resulting configuration is termed an individual network.

### 5.4. Computational Simulation and Logical Classification

Once the bZIP clusters were defined, each cluster was evaluated under a chemical-equilibrium framework to determine whether it could perform logical computation using EQTK. This Python library systematically calculates steady-state concentrations by solving the system’s reaction stoichiometry, experimental *K_a_* values, and mass conservation constraints. In each network configuration, two monomers were designated as inputs, one specific dimer acted as the output, and the remaining monomers served as accessory components.

Inspired by natural occurring expression levels, logical input states were defined by distinct concentrations: a logical zero (0) was represented by 10^−3^*nM* and a logical one (1) by 10^2^*nM* generating the four states of a two-input truth table. The upper concentration was motivated by cellular protein abundances on the order of tens of thousands of copies per cell^48,49^. The lower concentration was chosen to approximate the absence of the input species while its concentration in the equilibrium model remained nonzero.

Accessory monomer concentrations were allowed to vary within the same concentration interval. To identify the specific accessory monomer concentrations required to execute target logic gates, we implemented a heuristic optimization framework based on genetic algorithms. Conceptually, this process is analogous to training an artificial neural network, where the network architecture remains fixed while adjustable parameters are optimized to reproduce a desired input-output function. Here, however, the adjustable parameters are biochemical concentrations rather than numerical weights.

An individual network was classified as a successful logic gate if it maintained a strict, non-overlapping separation between its low and high concentration output states. Quantitatively, success required the maximum concentration recorded among all logical zeros (0) to be strictly lower than the minimum concentration recorded among all logical ones (1), this criterion ensures a non-overlapping threshold that is essential for robust downstream signal discrimination.

### 5.5. Optimization Framework via Genetic Algorithm

We implemented a customized genetic algorithm to identify the accessory monomer concentration vector 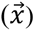 required for a cluster to compute a target Boolean logic gate while keeping experimental *K_a_* values fixed. Rather than the evaluation of continuous concentration titrations, each optimization run evaluated inputs as a discrete matrix of truth-table combinations, that corresponds to binary states (0 and 1). Meanwhile, accessory monomer concentrations were initialized randomly and iteratively refined by the algorithm.

Each candidate concentration vector was evaluated using EQTK to compute the equilibrium concentration of the candidate output dimer across all truth-table states. To measure performance, we calculated the Sum of Squared Errors (SSE) between the simulated equilibrium concentrations 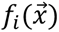 and the target concentration profiles *a_i_* that corresponds to the desired output logic states, as shown in **Eq. 4**. Target concentrations were assigned according to the desired logical outputs, using the same mapping as for the inputs: 10^−3^*nM* for logical zero and 10^2^*nM* for logical one.

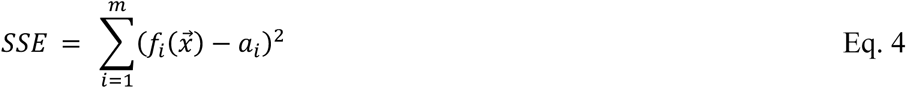

To guide the heuristic search, the SSE was mapped to a normalized fitness function (**Eq. 5**) with values from 0 (poor performance) to 1 (ideal, minimal-error solution). Through successive generations, the algorithm minimizes the discrepancy between simulated and target outputs and provides a reproducible mathematical process to identify concentration profiles capable of implementing target computational gates.

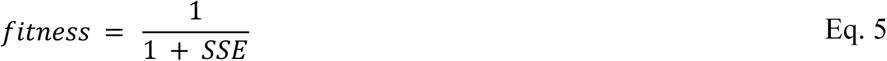

Optimization was performed using a real-coded genetic algorithm, where continuous values represented accessory monomer concentrations. Parent selection was conducted via binary tournament, and recombination was implemented using Simulated Binary Crossover (SBX)^58^ with a distribution index of *n_c_* = 5.

Optimization runs were executed for 500 generations with a population size of 300 individuals and a mutation rate parameter of 1. To ensure convergence and prevent loss of optimal solutions, an elitist strategy was applied by preserving the highest-fitness individual unchanged across generations.

### 5.6. Gate Evaluation Protocol for 2- and 3-Input Logic Operations

#### 5.6.1. Optimization of Accessory Concentrations for 2-Input Logic Gates and Computational Versatility

To quantify the computational capacity of each bZIP cluster under competitive dimerization, we first evaluated their performance across two-input Boolean operations. Input monomers were designated based on topological connectivity, prioritizing nodes with the highest degree of interaction, while all remaining monomers served as accessory species.

We measured the computational versatility of each cluster (defined as the number of unique two-input Boolean functions out of the 16 possible truth-table configurations) that it could execute solely by modulating accessory monomer concentrations.

All dimers formed within the cluster were considered as potential output species. In each optimization run, the equilibrium concentration profile of every candidate output dimer was compared against the desired truth-table output.

To establish computational versatility, optimized candidate solutions were primary-filtered using a high-accuracy fitness threshold (≥ 0.90). To avoid discarding functionally valid logic gates with lower raw fitness scores, we extended evaluation to solutions reaching a secondary threshold of ≥ 0.54. Empirical validation across our simulation trials confirmed that this lower boundary strictly maintains non-overlapping separation between logic-zero and logic-one output states, that guarantees robust downstream signal discrimination.

If a cluster achieved the defined fitness threshold for a given logic gate, it was classified as capable of computing that gate. In this case, the corresponding cluster, together with the optimized accessory monomer concentrations and selected output dimer, was considered an individual network.

#### 5.6.2. Scaling to Three-Input Logic Gates

To test whether these natural protein networks could support higher computational complexity, we selected the most versatile clusters identified in the two-input trials and subjected them to three-input logic gate optimization. We selected a benchmark suite of 17 foundational logic functions that represent different levels of computational complexity: AND, OR, MAJORITY (MAJ), MULTIPLEXER (MUX), XOR, NAND, NOR, XNOR, MINORITY (MIN), EXACTLY ONE, EXACTLY TWO, BUFFERS (A, B, and C), and INVERTERS (NOT A, NOT B, and NOT C).

Three-input configurations were evaluated under the same fitness thresholds as two-input gates to ensure strict, non-overlapping separation between logic-zero and logic-one concentration states.

### 5.7. Perturbation Analysis and Robustness to Expression Noise

Biological systems operate in inherently noisy intracellular environments where molecular abundances undergo continuous stochastic fluctuations^38^. To verify that our optimized bZIP networks maintain predictable Boolean logic under non-ideal biological conditions, we subjected each individual network to systematic concentration perturbations. This sensitivity analysis evaluates the operational boundaries and logical resilience of the computed gates across biologically relevant noise regimes.

#### 5.7.1. Stochastic Perturbation Framework

To simulate gene-expression variability, all molecular species were subjected to stochastic concentration perturbations drawn from a gamma distribution. This distribution was selected both to prevent unphysical negative concentrations and to reflect the underlying physics of transcriptional bursting in living cells, where steady-state protein abundances naturally follow gamma-distributed kinetics^59,60^.

To evaluate network stability across progressive noise intensities, we executed 1,000 independent stochastic perturbations for each optimized individual network across six target coefficients of variation (*CV* ∈ 0.05, 0.10, 0.20, 0.30, 0.40, 0.50). The perturbed concentration matrix (C_pert_) was defined as:

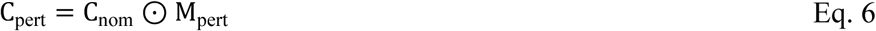

where *C_nom_* represents the baseline nominal concentrations, ⊙ denotes the element-wise Hadamard product, and *M*_pert_ is the random perturbation matrix. To ensure that the mean of the perturbed concentrations remains centered around the nominal baseline, the perturbation matrix is sampled from a gamma distribution with a mean (µ) equal to 1.0, while the standard deviation (σ) is set equal to the target coefficient of variation (*CV*), which specifies the desired percentage of noise error (using shape parameter *k* = 1/*CV*^2^ and scale parameter θ = *CV*^2^).

#### 5.7.2. Robustness Metrics and Performance Quantifiers

Robustness was evaluated applying three complementary metrics: probability of error, separation between logical states, and root mean squared deviation.

The probability of error (*P*_error_) was calculated as the ratio of misclassified perturbation trials to the total number of simulations (**Eq. 7**). A simulation was classified as a logical failure when stochastic perturbations induced an overlap between the output concentration distributions of logical zeros and logical ones, violating the nominal truth-table separation.

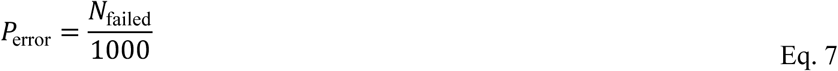

This metric indicates how frequently an individual network fails at a given level of extrinsic noise.

To evaluate the margin of discrimination between binary outputs under noise, we defined the signal separation (*S*, **Eq. 8**) across all 1,000 perturbation trials. This metric quantifies the distance between the 5th percentile of logic-one outputs (P_5_(Y_ones_)) and the 95th percentile of logic-zero outputs (*P*_95_(*Y*_zeros_)). Values of *S* > 1 indicate well-resolved, non-overlapping binary distributions, whereas *S* < 1 signifies output overlap. While *P*_error_ measures the absolute failure frequency of a gate, *S* captures the continuous degradation of its logical separation window.

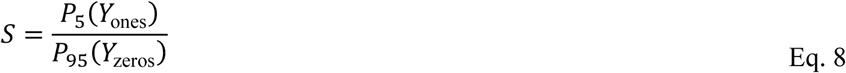

The third metric, the root mean squared deviation (*RMSD_lo_*_g_), quantifies how much the perturbed output profile deviates from the nominal output profile of a given network. For each simulation, *RMSD_lo_*_g_ was calculated across the *m* truth-table entries (**Eq. 9**), where *Output*_pert,*i*_ and *Output*_nom,*i*_ represent the perturbed and nominal output-dimer concentrations for the *i* − *th* input condition, respectively.

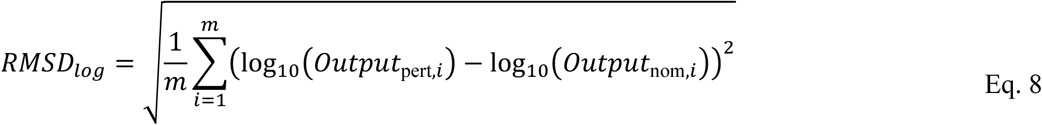

Across all 1,000 simulations per noise level, we evaluated the median *RMSD_lo_*_g_ along with its first and third quartiles (Q_25_ and Q_75_). The median *RMSD_lo_*_g_ describes the typical deformation of the output profile under noise, while the interquartile range (Q_75_ − Q_25_) reflects the response spread. A narrow interquartile range indicates that most perturbed outputs remain close to the nominal behavior and are not dominated by extreme perturbation events. Finally, to link these deviations to physical concentration changes, the overall nominal fold-change deviation can be estimated as 10^RMSD^*^log^*.

## Associated Content

### Supporting Information

The Supporting Information is available alongside the preprint main text on bioRxiv.

- Supplementary Figures S1–S21, Supplementary Tables S1–S8, and Supplementary Notes S1-S5 (PDF).
- Complete code and data will be available upon publication.

## Author Information

### Author Contributions

Conceptualization: EB, GMR, JAM. Experimental design: AOE, DFN, EB, GMR, JAM. Experimental execution: AOE, DFN. Formal analysis: AOE, DFN, EB, GMR, JAM Original draft preparation: AOE, DFN. Review and editing: AOE, DFN, GMR, EB, JAM

## Notes

The authors declare no competing financial interest.

## Acknowledgments

AOE and DFN acknowledge SECIHTI for scholarships Nos. 892202 and 2164900, respectively.

