## Supplementary Information for "Programmable Biochemical Logic Through Repurposing Natural bZIP Protein Interaction Networks"

### Supplementary Information for
### "Programmable Biochemical Logic Computation Through Repurposing Natural bZIP Protein Interaction Networks"

Arturo Orozco-Estrada[1], Daniela Flores-Nuño[1], Gerardo Mendizabal-Ruiz[1], Ernesto Borrayo[1,*], J. Alejandro Morales[1,*]

---

[1]Biodigital Innovation Lab, Translational Bioengineering Department, University of Guadalajara

Guadalajara 44430, Jalisco, Mexico

*

**This supplementary document contains:**

- Supplementary Figures S1–S2.
- Supplementary Tables S1–S8.
- Supplementary Notes S1-S5.

### Supplementary Note S1. Topological Network Analysis and Cluster Detection

This section presents the topological characterization of natural bZIP interaction networks and the identification of local communities as smaller substrates for computational analysis.

The bZIP interaction networks of *Nematostella vectensis*, *Ciona intestinalis*, and *Homo sapiens* were characterized using the experimental dataset reported by Reinke *et al.*[1] Curated interaction matrices are presented as heatmaps of dissociation constants ($K_d$), with yellow indicating weaker interactions and blue indicating stronger interactions. Network graphs represent proteins as nodes and pairwise interactions as edges weighted by association constants ($K_a$), with darker edges indicating stronger binding. Node colors distinguish the communities identified using the Louvain algorithm.

#### S1.1. Nematostella vectensis

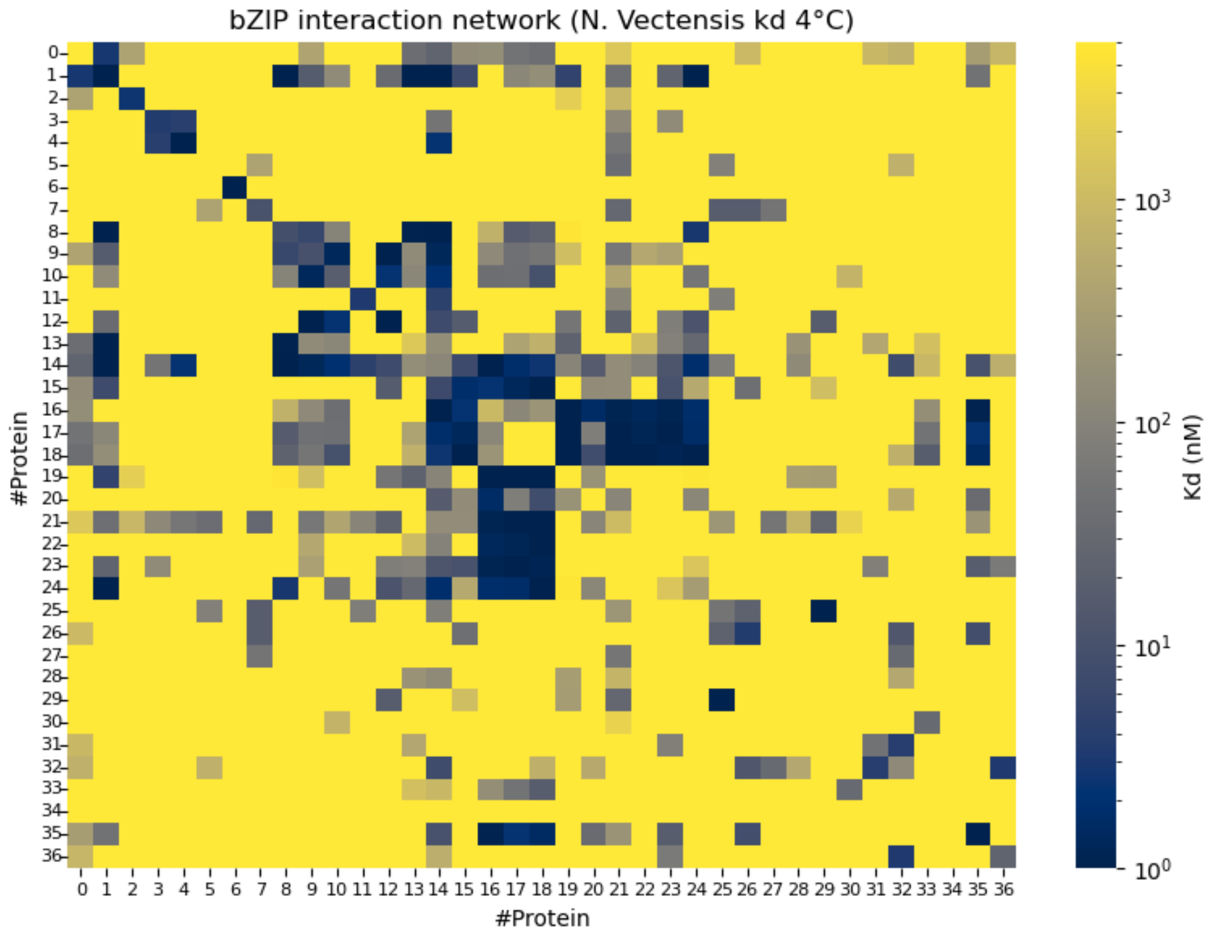


**Figure S1. Binding affinity matrix of the *Nematostella vectensis* bZIP network at 4 °C.**

Heatmap of curated pairwise dissociation constants $K_d$ $(nM)$ for 37 bZIP proteins from *Nematostella vectensis*, based on experimental data reported by Reinke *et al.*[1] Rows and columns correspond to individual proteins, indexed from 0 to 36. The logarithmic color scale ranges from blue, indicating stronger interactions (lower $K_d$), to yellow, indicating weaker interactions (higher $K_d$). Diagonal entries represent homodimeric interactions, whereas off-diagonal entries represent heterodimeric interactions. The curated network contains 196 interactions among 703 possible unique protein pairs, including homodimers.

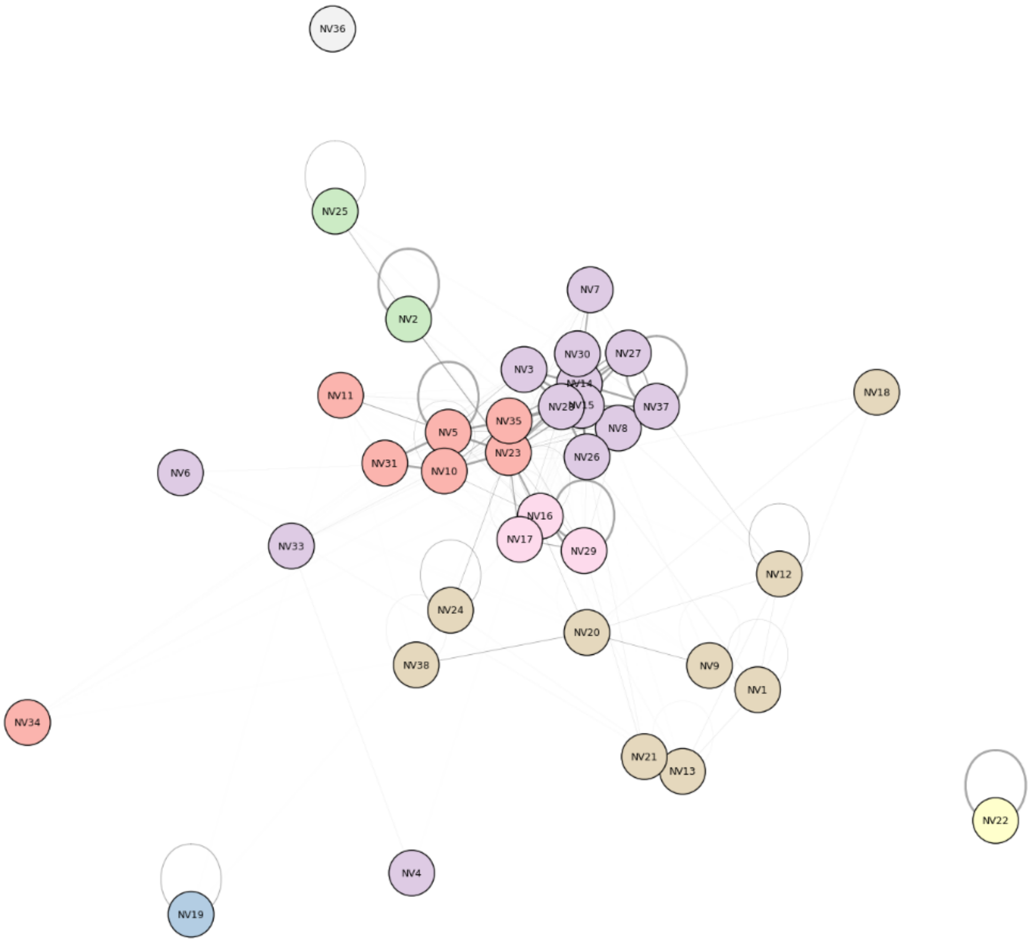


**Figure S2. Graph representation and community structure of the *Nematostella vectensis* bZIP interaction network at 4 °C.**

Graph representation of the curated *Nematostella vectensis* bZIP interaction network. Nodes represent individual proteins, and edges represent pairwise interactions weighted by association constants ($K_a$), with darker edges indicating stronger interactions. Self-loops denote homodimerization. Node colors distinguish the eight communities identified using the Louvain algorithm, ranging in size from single proteins to a cluster containing 13 proteins. Most proteins belong to a large connected component, with edges linking different communities, while a few proteins remain disconnected from the rest of the network.

**Table S1. Structural properties of *Nematostella vectensis* bZIP clusters selected for logic gate optimization at 4°C.**

The table summarizes the composition and connectivity of the selected clusters. For each cluster, "Cluster ID" indicates its original label in the Louvain partition of the complete network. "Number of proteins" and "Included proteins" report the cluster size and the identities of its constituent proteins, respectively. "Number of interactions" indicates how many active protein pairs interact within the cluster, including both homodimeric and heterodimeric interactions. "Total possible pairs" represents all unique homodimeric and heterodimeric combinations among the proteins in the cluster, regardless of whether an interaction was recorded.

| *N. vectensis* | | | | |
|---|---|---|---|---|
| **ID CLUSTER** | **Number of proteins** | **Included proteins** | **Number of interactions** | **Total possible pairs** |
| #0 | 7 | 'NV11', 'NV5', 'NV10', 'NV31', 'NV23', 'NV35', 'NV34' | 20 | 28 |
| #3 | 13 | 'NV6', 'NV8', 'NV14', 'NV15', 'NV28', 'NV3', 'NV7', 'NV26', 'NV27', 'NV30', 'NV4', 'NV33', 'NV37' | 41 | 91 |
| #5 | 9 | 'NV1', 'NV24', 'NV13', 'NV12', 'NV18', 'NV21', 'NV9', 'NV20', 'NV38' | 17 | 45 |

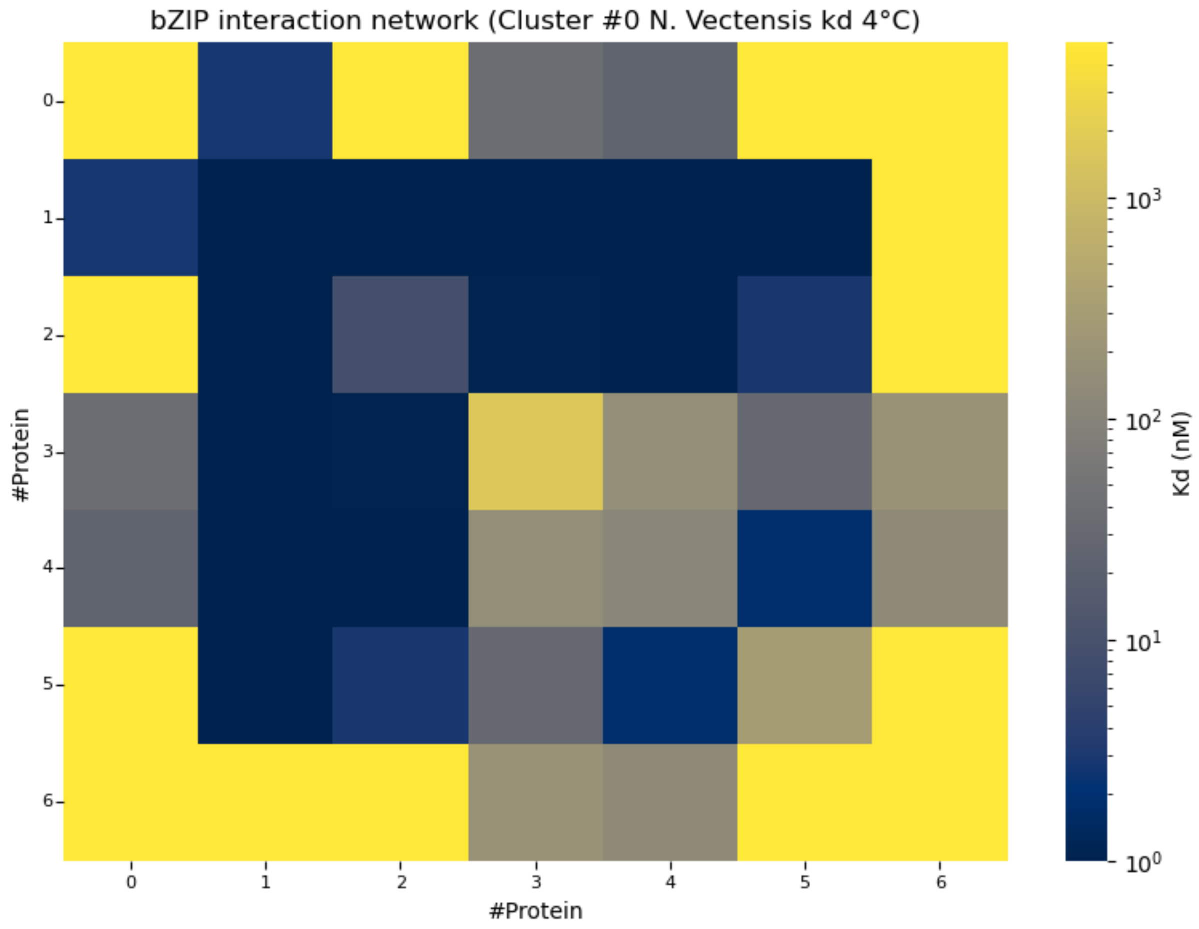


**Figure S3. Binding affinity matrix of *Nematostella vectensis* bZIP cluster #0 at 4°C.**

Pairwise binding affinities among the seven bZIP proteins comprising *Nematostella vectensis* cluster #0 at 4°C. Dissociation constants $K_d$ $(nM)$ are displayed on a logarithmic color scale, with blue indicating stronger interactions and yellow indicating weaker interactions. This cluster contains 20 of the 28 possible unique homodimeric and heterodimeric interactions, yielding a connectivity density of 71.4%, the highest among the selected clusters from this species. Rows and columns are indexed from zero according to the protein order in **Table S1**. NV31 and NV23, corresponding to indices 3 and 4, respectively, are the most highly connected proteins within the cluster.

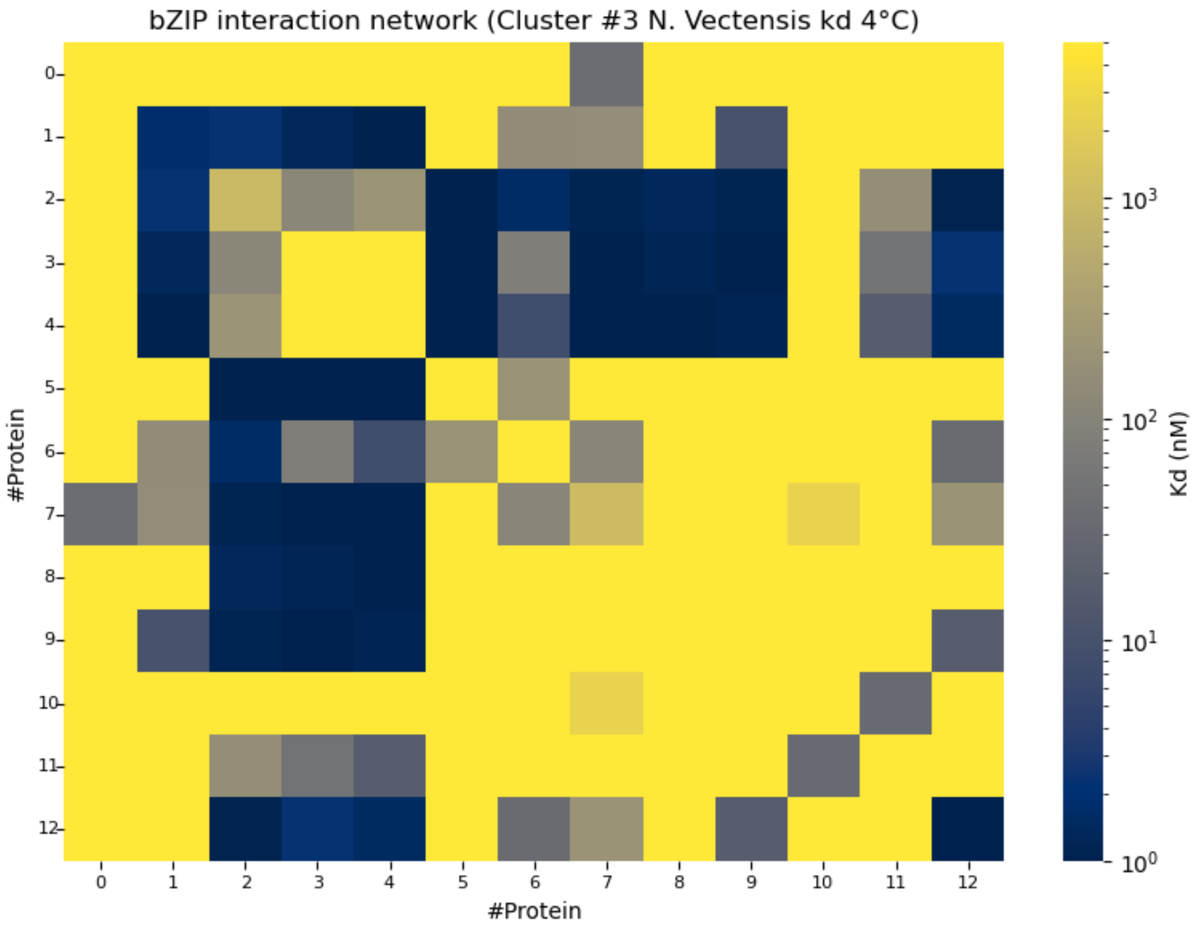


**Figure S4. Binding affinity matrix of *Nematostella vectensis* bZIP cluster #3 at 4°C.**

Cluster #3 comprises 13 bZIP proteins and has the greatest number of internal interactions among the selected *Nematostella vectensis* clusters. Its 41 interactions account for 45.1% of the 91 possible unique dimer pairs. The matrix displays dissociation constants $K_d$ $(nM)$ on a logarithmic scale, ranging from blue for stronger binding to yellow for weaker binding. Rows and columns are indexed from zero according to the protein order in **Table S1**. NV14 and NV28, labeled 2 and 4, respectively, exhibit the highest connectivity.

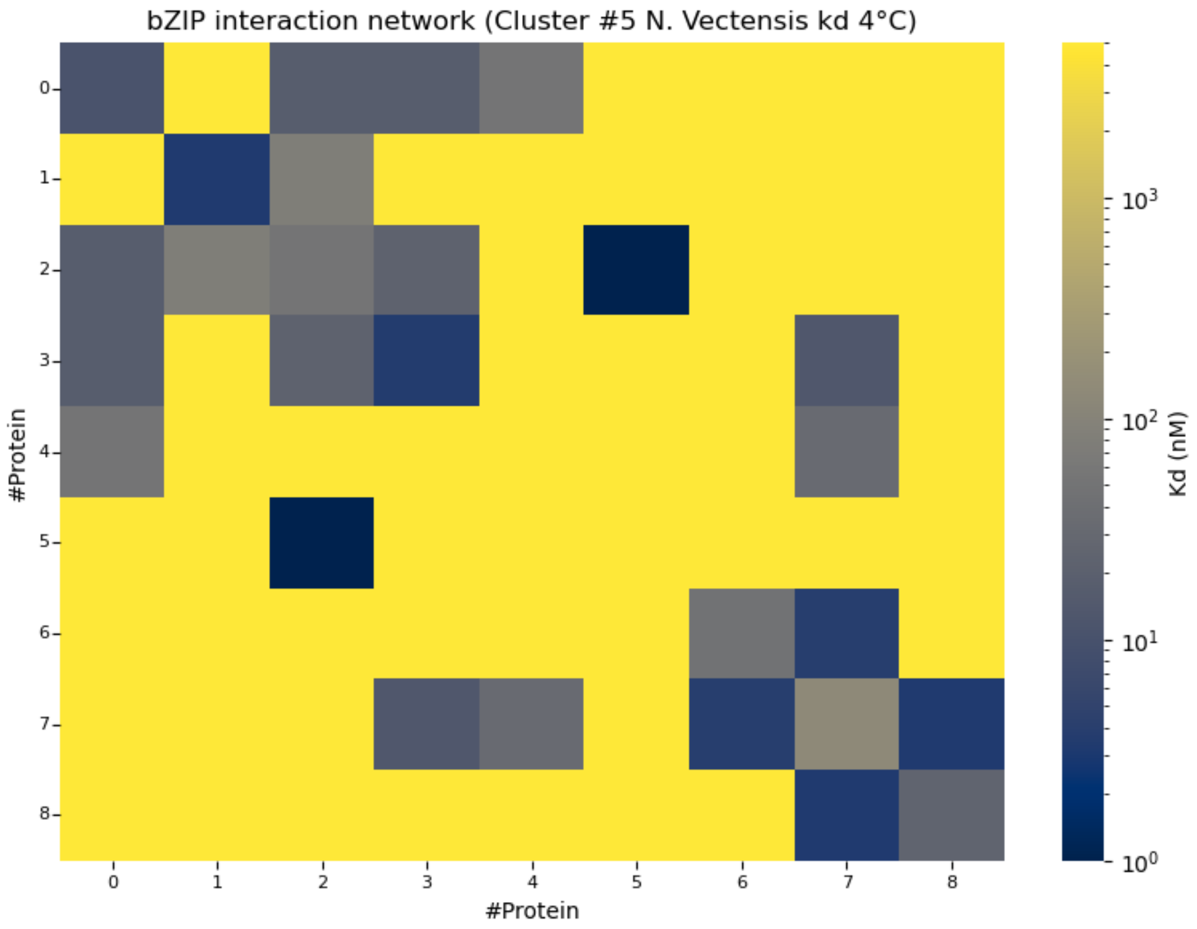


**Figure S5. Binding affinity matrix of *Nematostella vectensis* bZIP cluster #5 at 4°C.**

The nine proteins in cluster #5 form a network with 17 interactions among 45 possible unique dimer pairs. Its connectivity density of 37.8% is the lowest among the selected *Nematostella vectensis* clusters. Matrix entries represent dissociation constants $K_d$ $(nM)$ measured at 4°C, with a logarithmic color scale distinguishing stronger interactions in blue from weaker interactions in yellow. Rows and columns are indexed from zero according to the protein order in **Table S1**. NV13 and NV20, corresponding to indices 2 and 7, respectively, are the most highly connected proteins.

#### S1.2. *Ciona intestinalis*

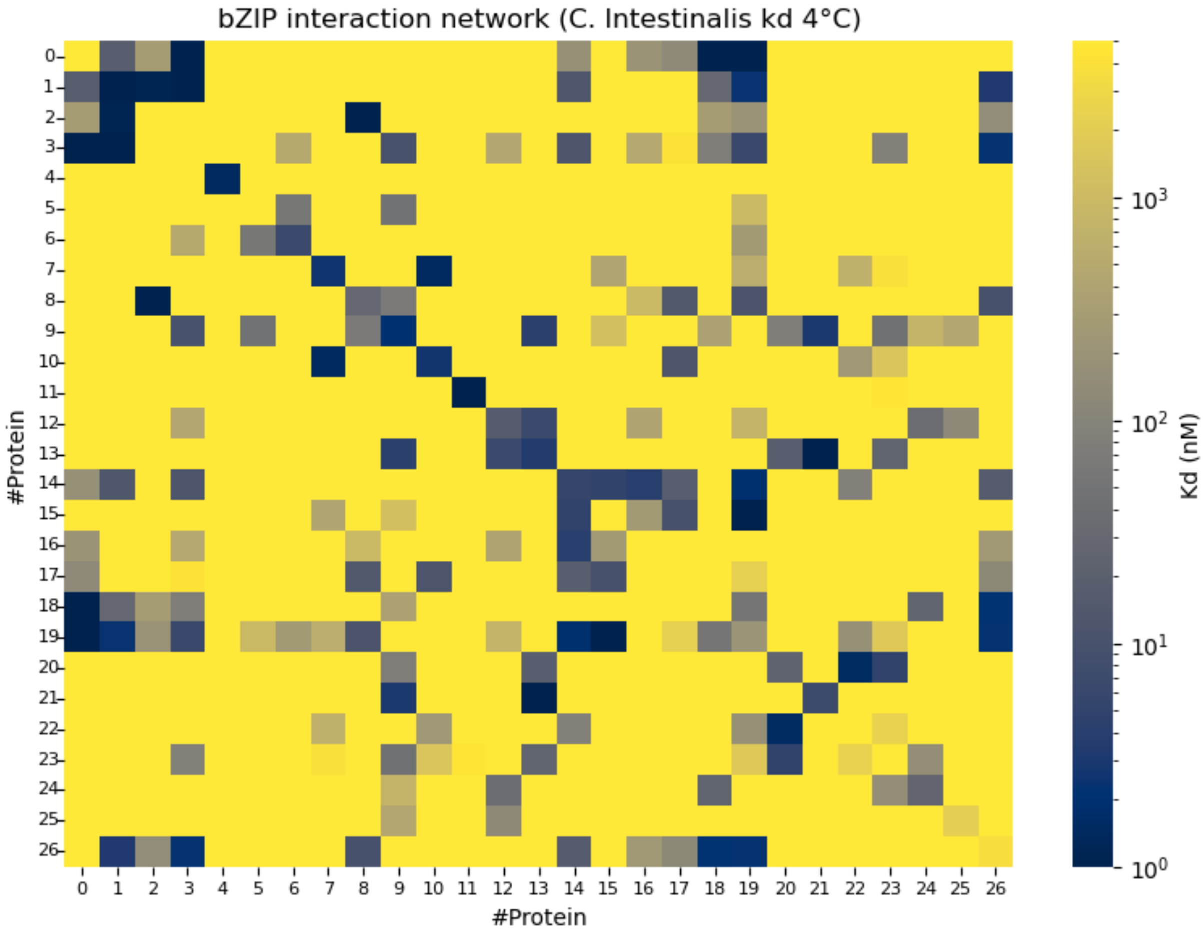


**Figure S6. Binding affinity matrix of the *Ciona intestinalis* bZIP network at 4°C.**

The matrix summarizes pairwise binding affinities among the 27 bZIP proteins in the curated *Ciona intestinalis* network, comprising 101 interactions among 378 possible unique dimer pairs. Dissociation constants $K_d$ $(nM)$ at 4°C are displayed on a logarithmic color scale, with blue indicating stronger interactions and yellow indicating weaker interactions. In the displayed protein ordering, high-affinity interactions appear as scattered, localized patches separated by regions of weaker binding. This fragmented pattern suggests that strong interactions are concentrated within several local groups.

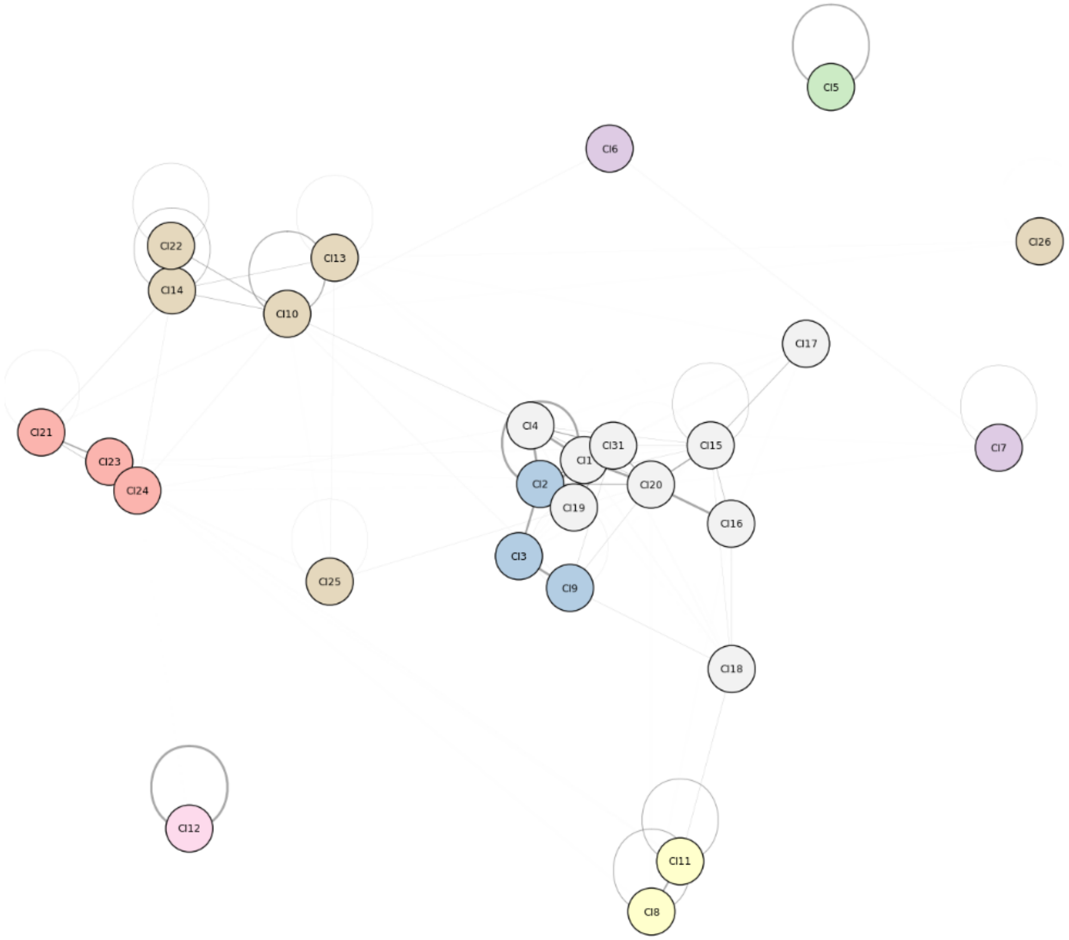


**Figure S7. Graph representation and community structure of the *Ciona intestinalis* bZIP interaction network at 4°C.**

Graph representation of the *Ciona intestinalis* bZIP interaction network. Nodes represent individual bZIP proteins, and edges denote pairwise interactions weighted by association constants ($K_a$), with darker edges indicating stronger binding. Self-loops represent homodimerization, and node colors distinguish the eight communities identified by the Louvain algorithm, ranging from single proteins to a cluster of nine proteins. The *Ciona intestinalis* graph exhibits more separated communities, with sparse and predominantly weak interactions between clusters. This organization suggests limited coupling between communities within the network.

**Table S2. Structural properties of *Ciona intestinalis* bZIP clusters selected for logic gate optimization at 4°C.**

The table summarizes the composition and connectivity of clusters #0, #1, #5, and #7 selected from the *Ciona intestinalis* network for biochemical simulations. For each cluster, the number and names of its constituent proteins are provided alongside the number of unique interactions, including homodimeric and heterodimeric pairs, each counted once. The total possible pairs include all unique dimer combinations among the cluster's proteins, regardless of whether an interaction was recorded.

| C. Intestinalis | | | | |
|---|---|---|---|---|
| **ID CLUSTER** | **Number of proteins** | **Included proteins** | **Number of interactions** | **Total possible pairs** |
| #0 | 3 | 'CI21', 'CI23', 'CI24' | 4 | 6 |
| #1 | 3 | 'CI2', 'CI3', 'CI9' | 4 | 6 |
| #5 | 6 | 'CI10', 'CI13', 'CI14', 'CI22', 'CI25', 'CI26' | 14 | 21 |
| #7 | 9 | 'CI1', 'CI4', 'CI15', 'CI16', 'CI17', 'CI18', 'CI19', 'CI20', 'CI31' | 29 | 45 |

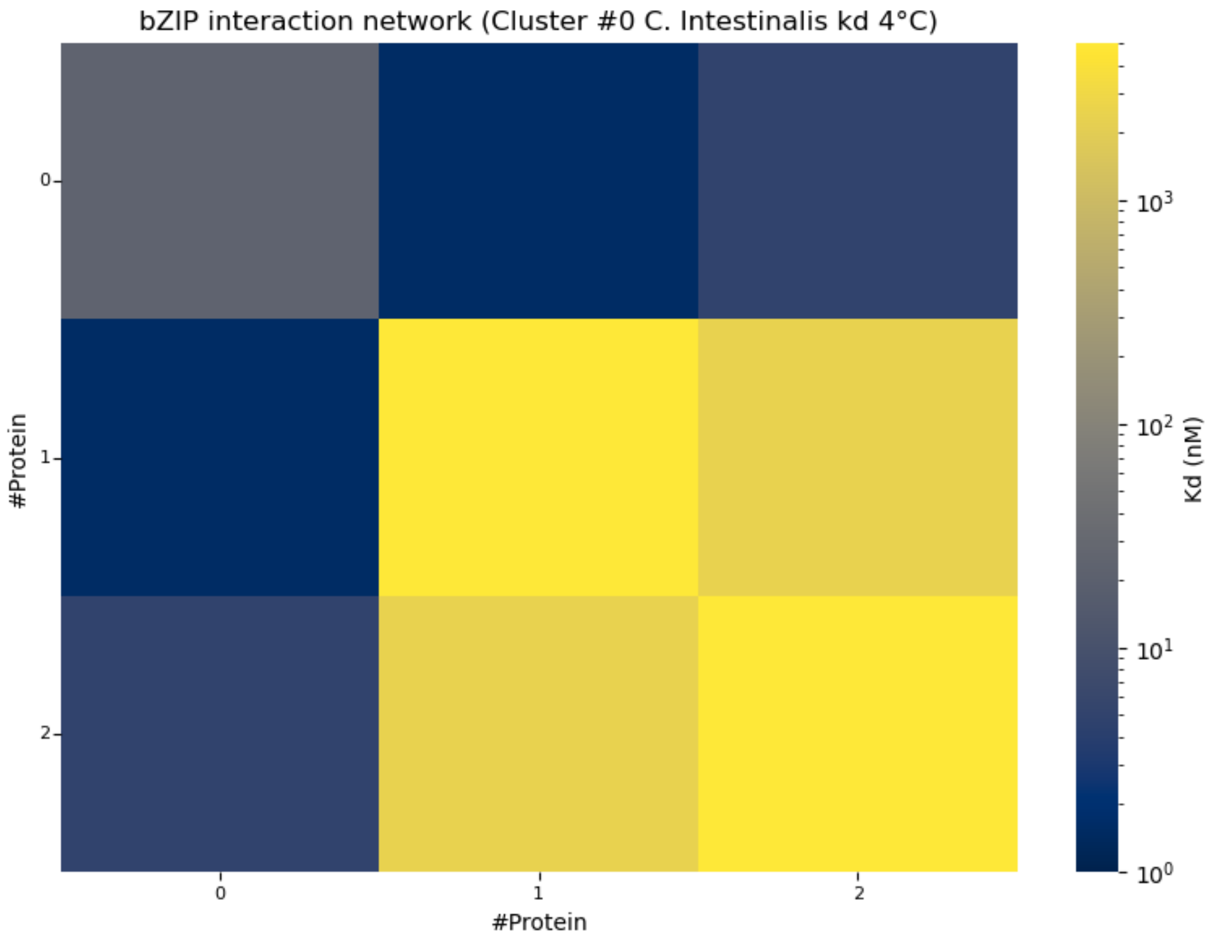


**Figure S8. Binding affinity matrix of *Ciona intestinalis* bZIP cluster #0 at 4°C.**

Cluster #0 comprises three bZIP proteins CI21, CI23, and CI24 and contains four interactions among six possible unique dimer pairs, corresponding to a connectivity density of 66.7%. The heatmap displays dissociation constants $K_d$ $(nM)$ on a logarithmic color scale, with blue indicating stronger interactions and yellow indicating weaker interactions. Protein indices follow the order listed in **Table S2**, starting from zero. CI21 and CI23, corresponding to indices 0 and 1, respectively, are the most highly connected proteins within the cluster.

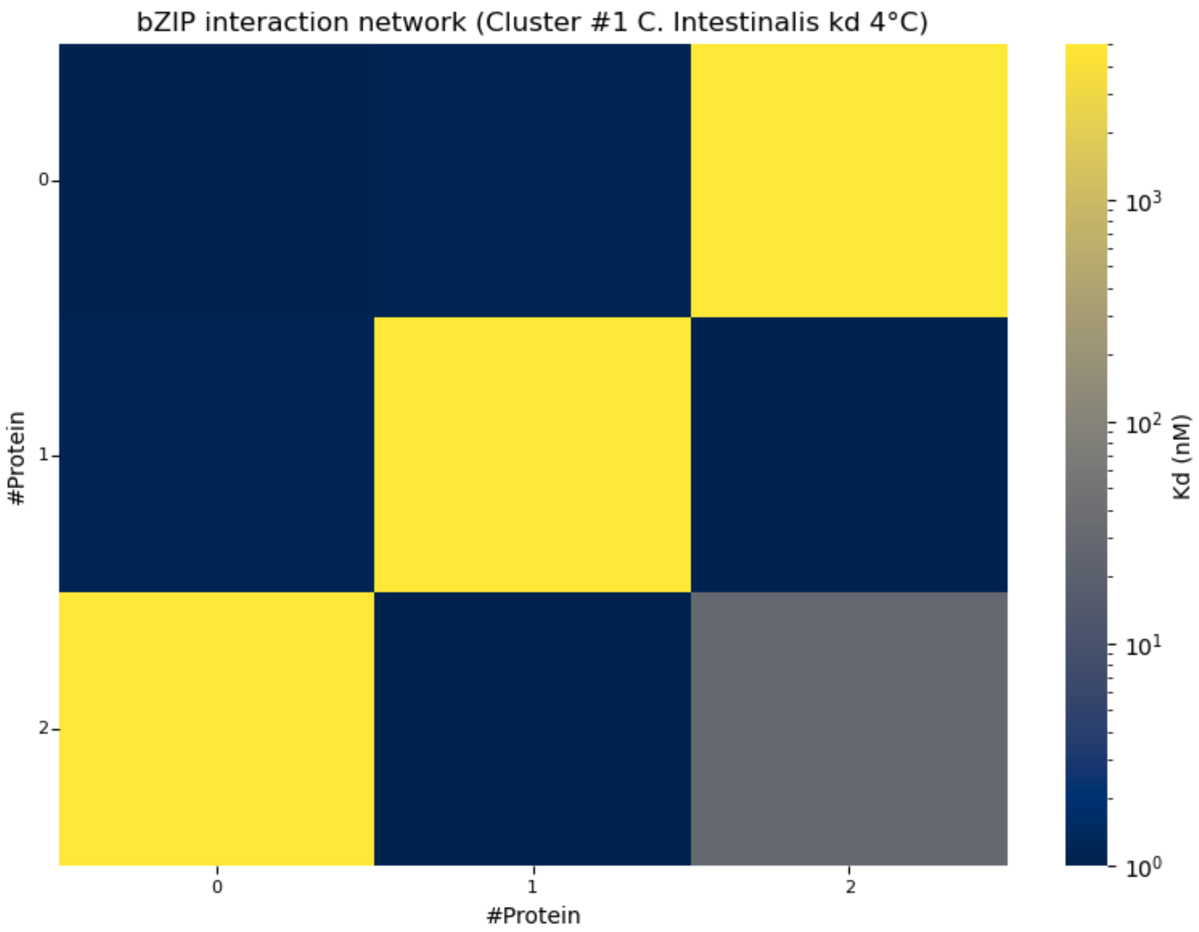


**Figure S9. Binding affinity matrix of *Ciona intestinalis* bZIP cluster #1 at 4°C.**

Cluster #1 consists of CI2, CI3, and CI9, with four interactions among six possible unique dimer pairs. Like cluster #0 (**Figure S8**), it has a connectivity density of 66.7% and provides a minimal configuration for two input monomers and one accessory monomer. The heatmap displays dissociation constants $K_d$ $(nM)$ on a logarithmic color scale, with blue indicating stronger interactions and yellow indicating weaker interactions. Protein indices follow the order listed in **Table S2**, starting from zero. CI2 and CI3, indexed as 0 and 1, respectively, have the highest connectivity within the cluster.

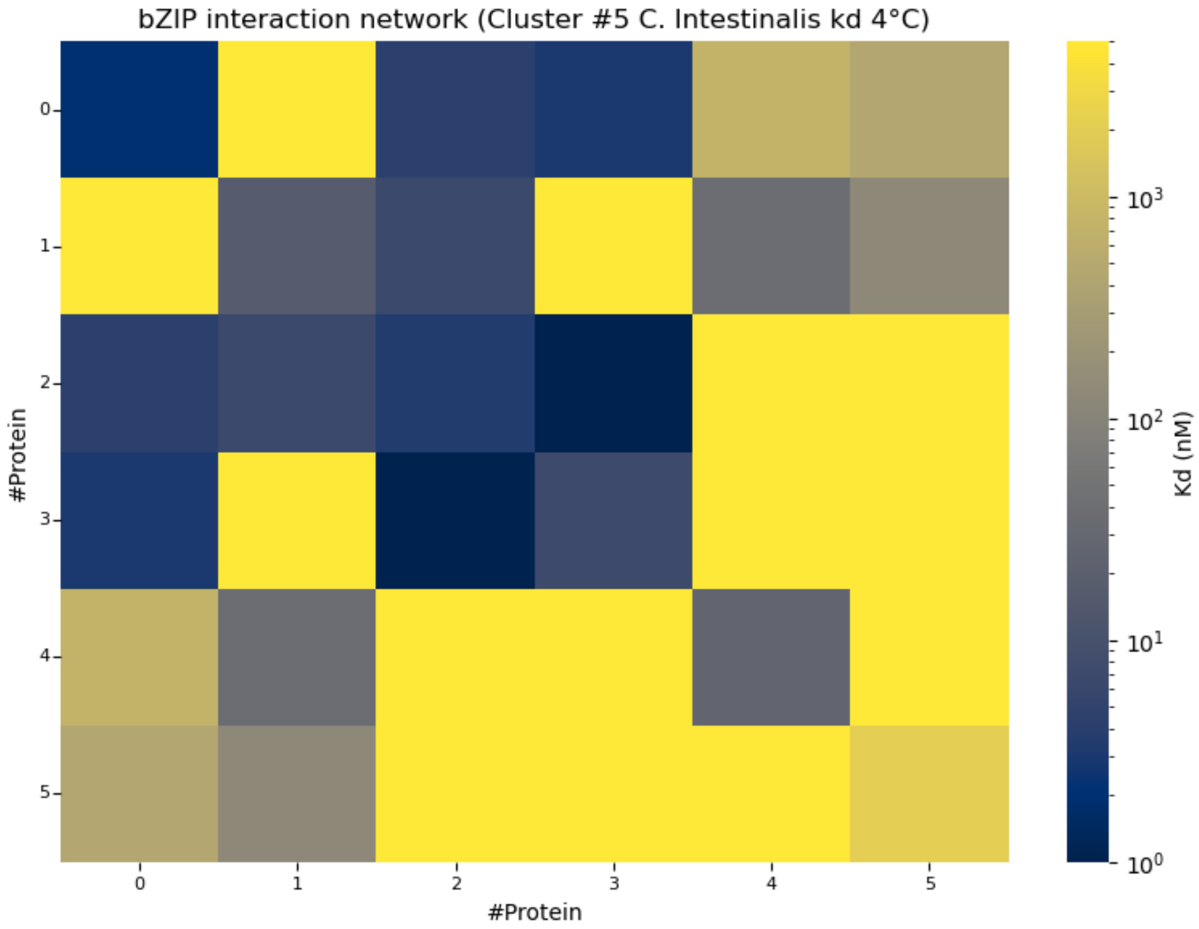


**Figure S10. Binding affinity matrix of *Ciona intestinalis* bZIP cluster #5 at 4 °C.**

Cluster #5 comprises six bZIP proteins connected by 14 interactions among 21 possible unique dimer pairs. Its connectivity density of 66.7% matches that of clusters #0 and #1 despite its larger size. The heatmap presents dissociation constants $K_d$ $(nM)$ on a logarithmic scale, with blue denoting stronger binding and yellow denoting weaker binding. Protein indices follow the order listed in **Table S2**, starting from zero. CI10 and CI14, corresponding to indices 0 and 2, respectively, are the most highly connected proteins within the cluster.

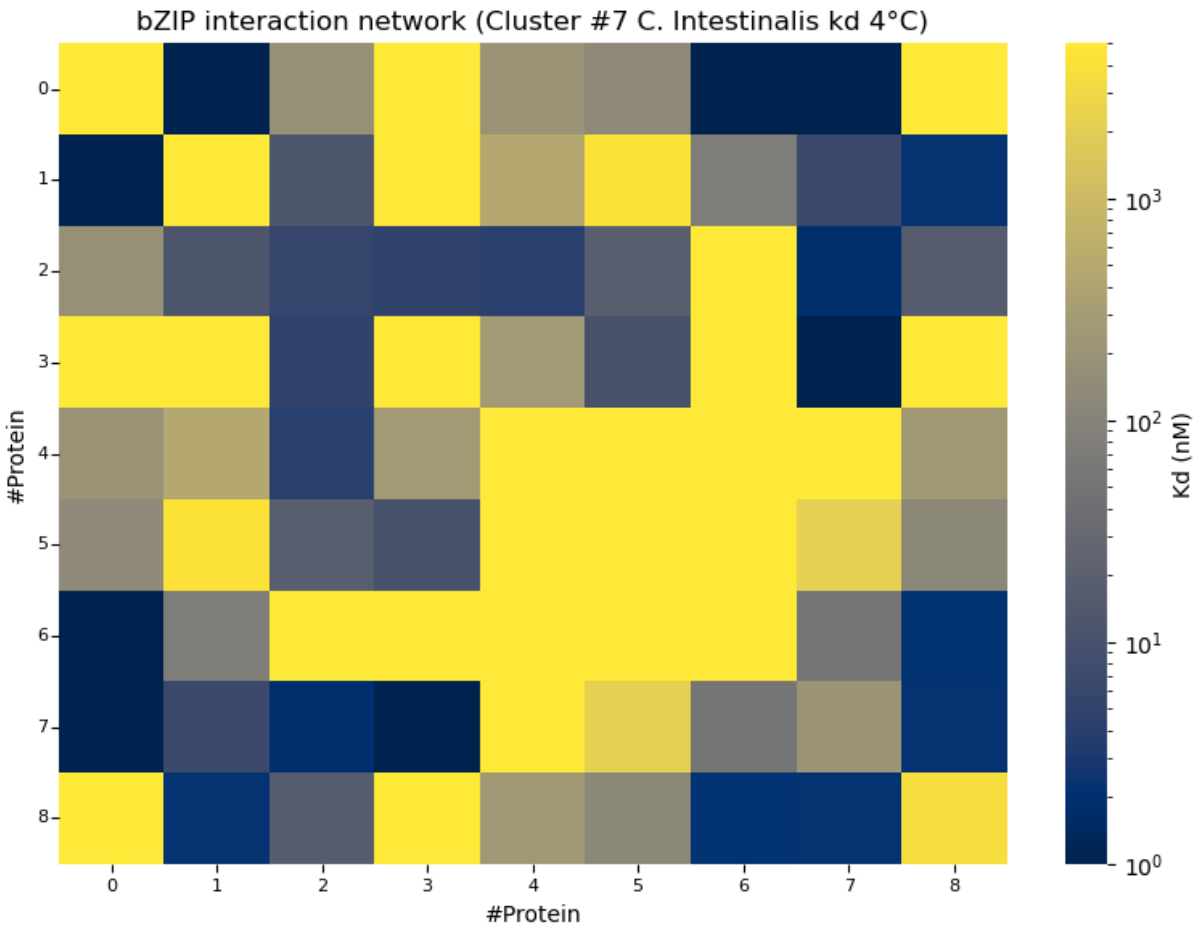


**Figure S11. Binding affinity matrix of *Ciona intestinalis* bZIP cluster #7 at 4°C.**

Cluster #7 is the largest of the selected *Ciona intestinalis* communities, comprising nine bZIP proteins and 29 interactions among 45 possible unique dimer pairs. Its connectivity density of 64.4% is slightly lower than that of the other selected clusters. Matrix entries show dissociation constants $K_d$ $(nM)$ on a logarithmic color scale, with stronger interactions represented in blue and weaker interactions in yellow. Protein indices follow the order listed in **Table S2**, starting from zero. CI20 and CI15, indexed as 7 and 2, respectively, exhibit the highest connectivity within the cluster.

#### S1.3. *Homo sapiens*

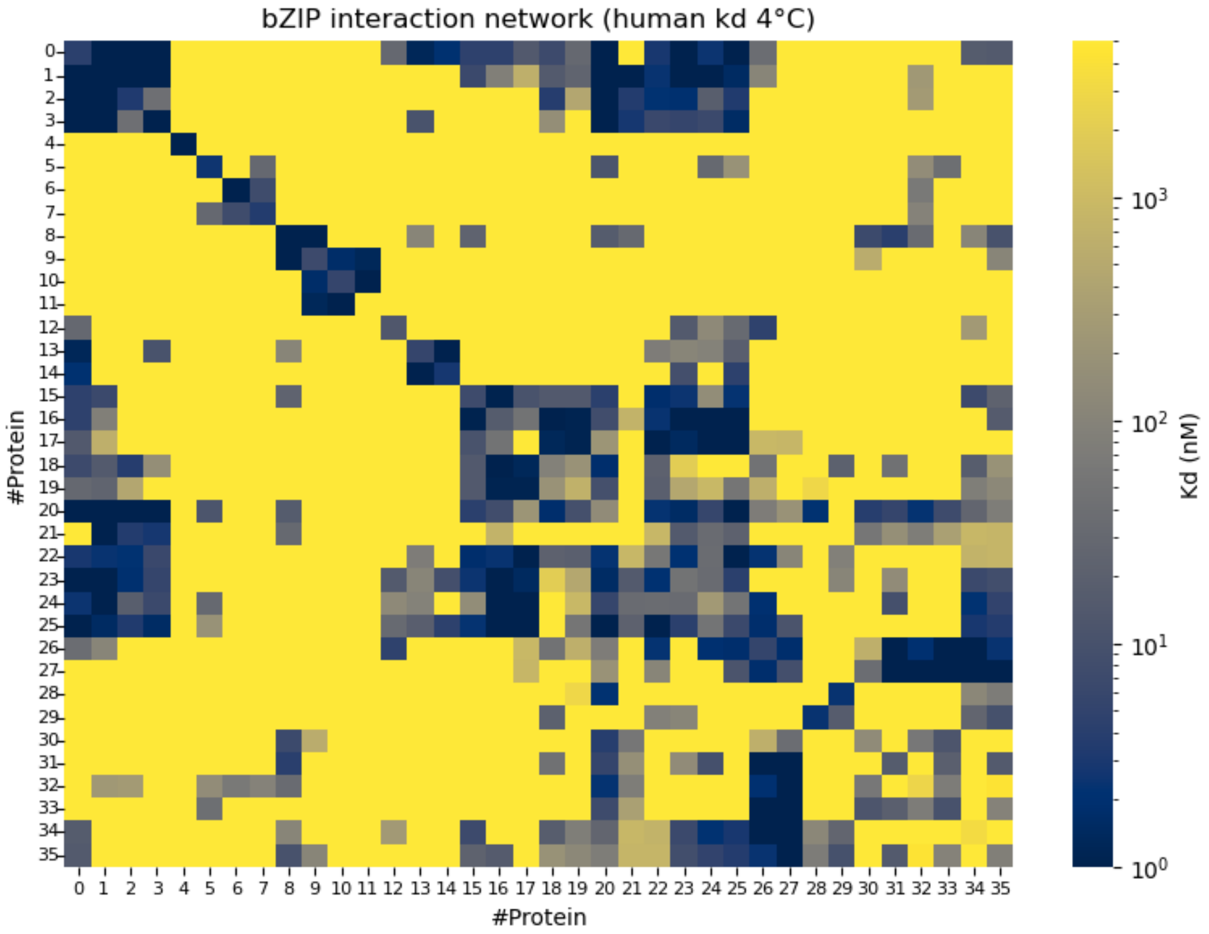


**Figure S12. Binding affinity matrix of the *Homo sapiens* bZIP network at 4°C.**

The matrix presents pairwise binding affinities among 36 bZIP proteins in the curated *Homo sapiens* network at 4°C. This network contains 239 interactions among 666 possible unique homodimeric and heterodimeric pairs, corresponding to a connectivity density of 35.9%, the highest among the three complete species networks analyzed at this temperature. Rows and columns represent individual proteins, and matrix entries show dissociation constants $K_d$ $(nM)$ on a logarithmic color scale, with blue indicating stronger interactions and yellow indicating weaker interactions.

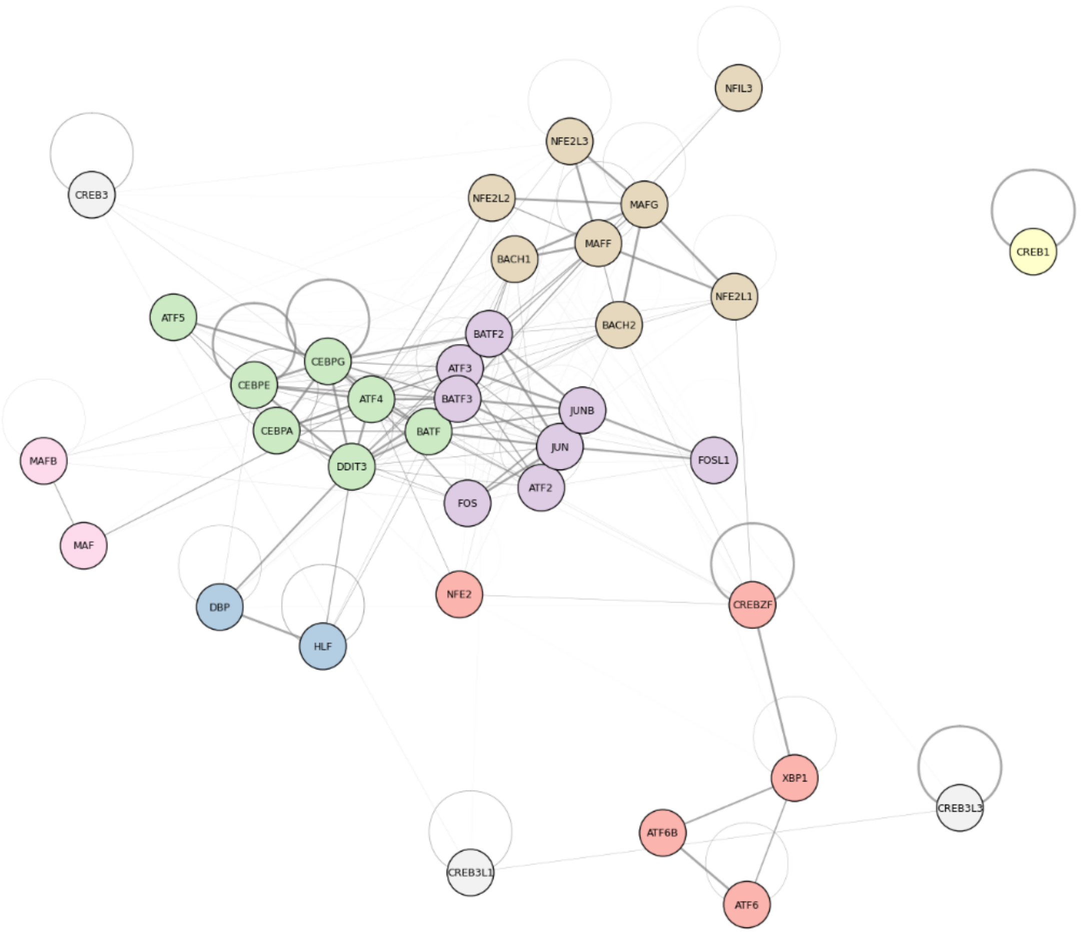


**Figure S13. Graph representation and community structure of the *Homo sapiens* bZIP interaction network at 4°C.**

The graph represents interactions among the 36 bZIP proteins in the *Homo sapiens* network at 4°C. Nodes correspond to individual proteins, and edges represent pairwise interactions weighted by association constants $K_a$, with darker edges indicating stronger binding. Self-loops denote homodimerization. Node colors distinguish the eight communities identified by the Louvain algorithm, ranging from single-protein communities to two clusters containing eight proteins each. Connections between communities illustrate that the detected clusters remain linked within the broader interaction network.

**Table S3. Structural properties of *Homo sapiens* bZIP clusters selected for logic gate optimization at 4°C.**

The table summarizes the composition and connectivity of the five selected *Homo sapiens* clusters: #0, #2, #3, #5, and #7. The number and names of its constituent proteins describe its size and composition, while the number of interactions indicates the active homodimeric and heterodimeric pairs present within the cluster. The total possible pairs include all unique dimer combinations among these proteins, whether or not an interaction is present in the network.

| ***Homo Sapiens*** | | | | |
|---|---|---|---|---|
| **ID CLUSTER** | **Number of proteins** | **Included proteins** | **Number of interactions** | **Total possible pairs** |
| #0 | 5 | 'CREBZF', 'XBP1', 'ATF6', 'ATF6B', 'NFE2' | 10 | 15 |
| #2 | 8 | 'ATF2', 'JUN', 'JUNB', 'FOS', 'FOSL1', 'ATF3', 'BATF2', 'BATF3' | 33 | 36 |
| #3 | 7 | 'DDIT3', 'CEBPG', 'CEBPA', 'CEBPE', 'ATF4', 'ATF5', 'BATF' | 25 | 28 |
| #5 | 8 | 'NFIL3', 'MAFF', 'MAFG', 'NFE2L1', 'NFE2L2', 'NFE2L3', 'BACH1', 'BACH2' | 26 | 36 |
| #7 | 3 | 'CREB3', 'CREB3L3', 'CREB3L1' | 5 | 6 |

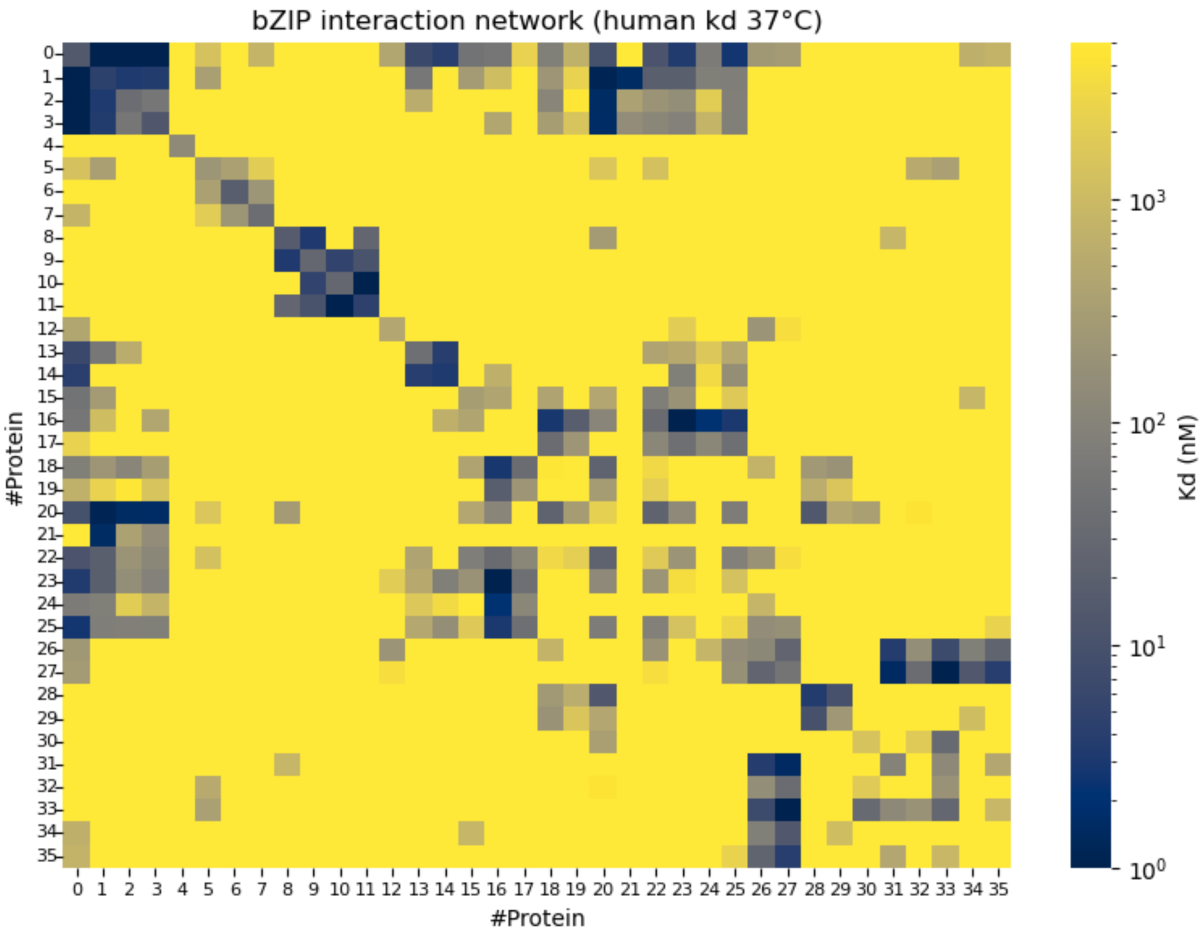


**Figure S14. Binding affinity matrix of the *Homo sapiens* bZIP network at 37°C.**

The matrix displays pairwise dissociation constants $K_d\ (nM)$ for the 36 proteins in the curated *Homo sapiens* bZIP network at 37°C. Colors follow a logarithmic scale, with blue indicating stronger interactions and yellow indicating weaker interactions. Four localized regions of stronger binding remain apparent, broadly resembling the pattern observed at 4°C (**Figure S12**). The overall shift toward weaker binding is accompanied by a reduction in the number of interactions from 239 at 4° to 173 at 37°C, among 666 possible unique dimer pairs, corresponding to a connectivity density of 26.0%.

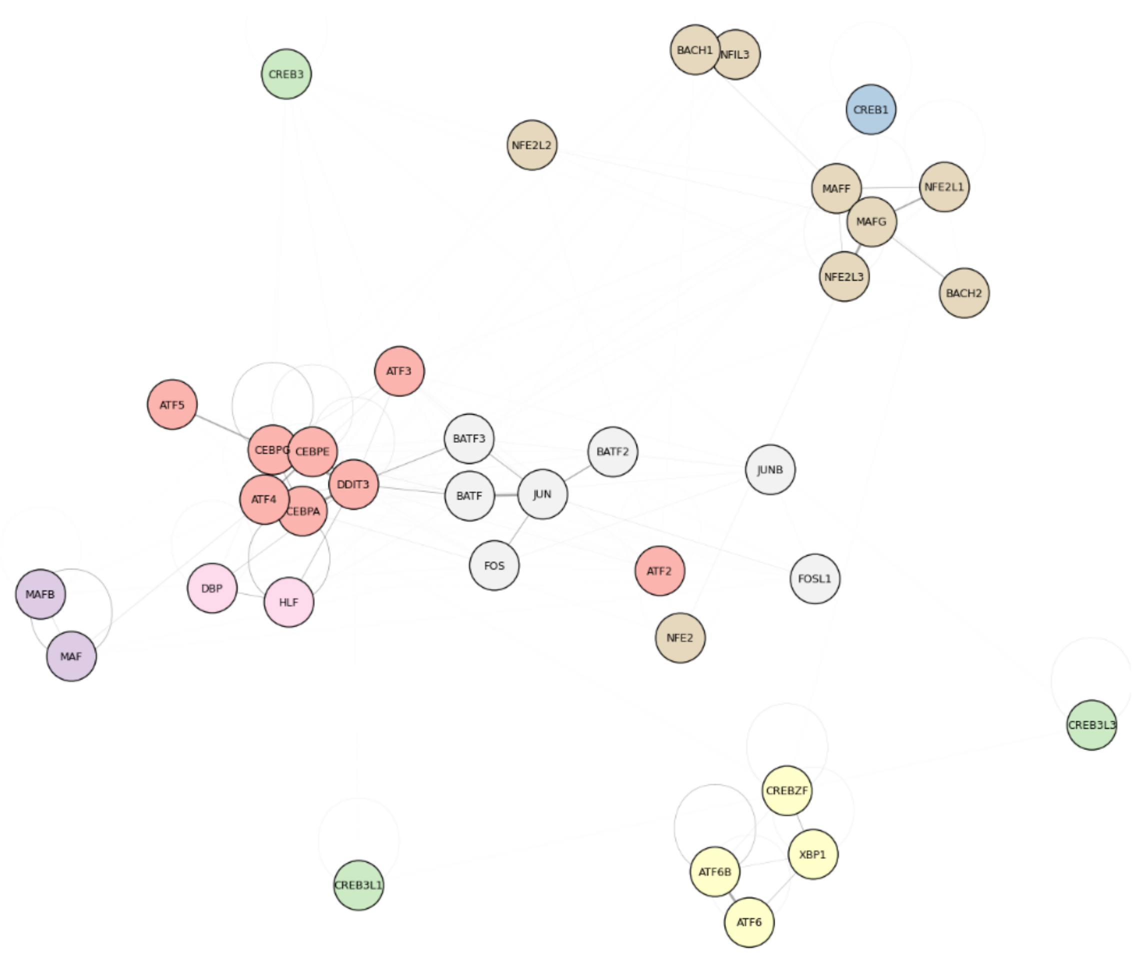


**Figure S15. Graph representation and community structure of the *Homo sapiens* bZIP interaction network at 37°C.**

The graph shows the community organization of the *Homo sapiens* bZIP interaction network at 37°C, with fewer connections between communities than observed at 4°C (**Figure S13**). The Louvain algorithm identifies eight communities at both temperatures, although their protein composition differs. Nodes represent individual proteins, and their colors indicate community membership. Edges represent pairwise interactions weighted by association constants $K_a$, with darker edges indicating stronger binding and self-loops denoting homodimerization. The reduced inter-community connectivity illustrates a more compartmentalized network topology at 37°C.

**Table S4. Structural properties of *Homo sapiens* bZIP cluster #0 selected for logic gate optimization at 37°C.**

The table summarizes the composition and connectivity of Homo sapiens bZIP cluster #0 at 37°C, which corresponds to the community represented by cluster #3 at 4°C. At 37°C, its composition changes and its size increases from seven to eight proteins. The cluster contains 29 interactions among 36 possible unique dimer pairs. Protein names indicate its composition, while the interaction count includes each homodimeric and heterodimeric pair present in the curated network. The total possible pairs include all unique dimer combinations among its proteins.

| ***Homo Sapiens* 37°C** | | | | |
|---|---|---|---|---|
| **ID CLUSTER** | **Number of proteins** | **Included proteins** | **Number of interactions** | **Total possible pairs** |
| #0 | 8 | 'DDIT3', 'CEBPG', 'CEBPA', 'CEBPE', 'ATF2', 'ATF4', 'ATF5', 'ATF3' | 29 | 36 |

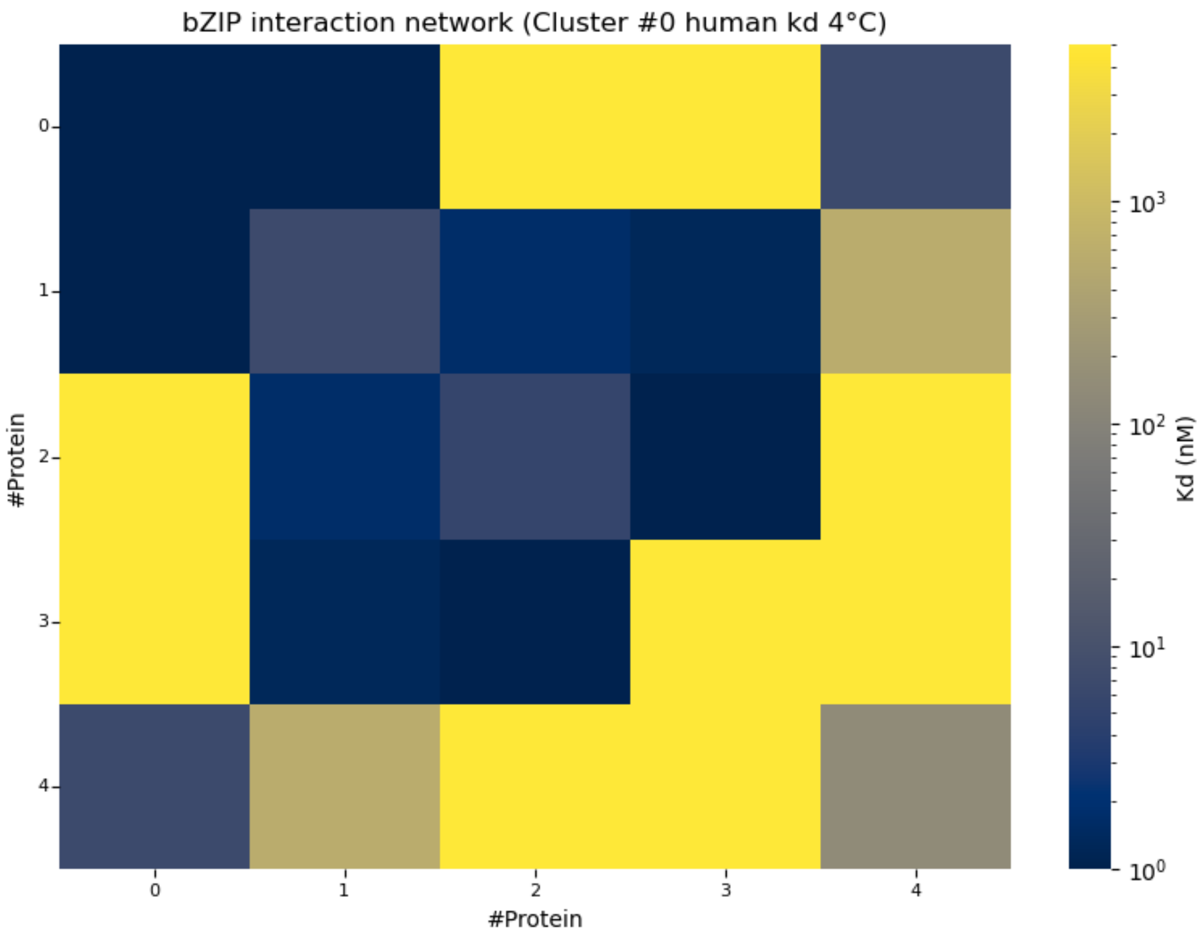


**Figure S16. Binding affinity matrix of *Homo sapiens* bZIP cluster #0 at 4°C.**

The matrix displays pairwise binding affinities among the five proteins comprising *Homo sapiens* cluster #0 at 4°C. This cluster contains 10 interactions among 15 possible unique dimer pairs, corresponding to a connectivity density of 66.7%. Dissociation constants $K_d$ $(nM)$ are shown on a logarithmic color scale. Protein indices start at 0 and follow the order listed for this cluster in **Table S3**. CREBZF and XBP1, indexed as 0 and 1, respectively, are the most highly connected proteins within the cluster.

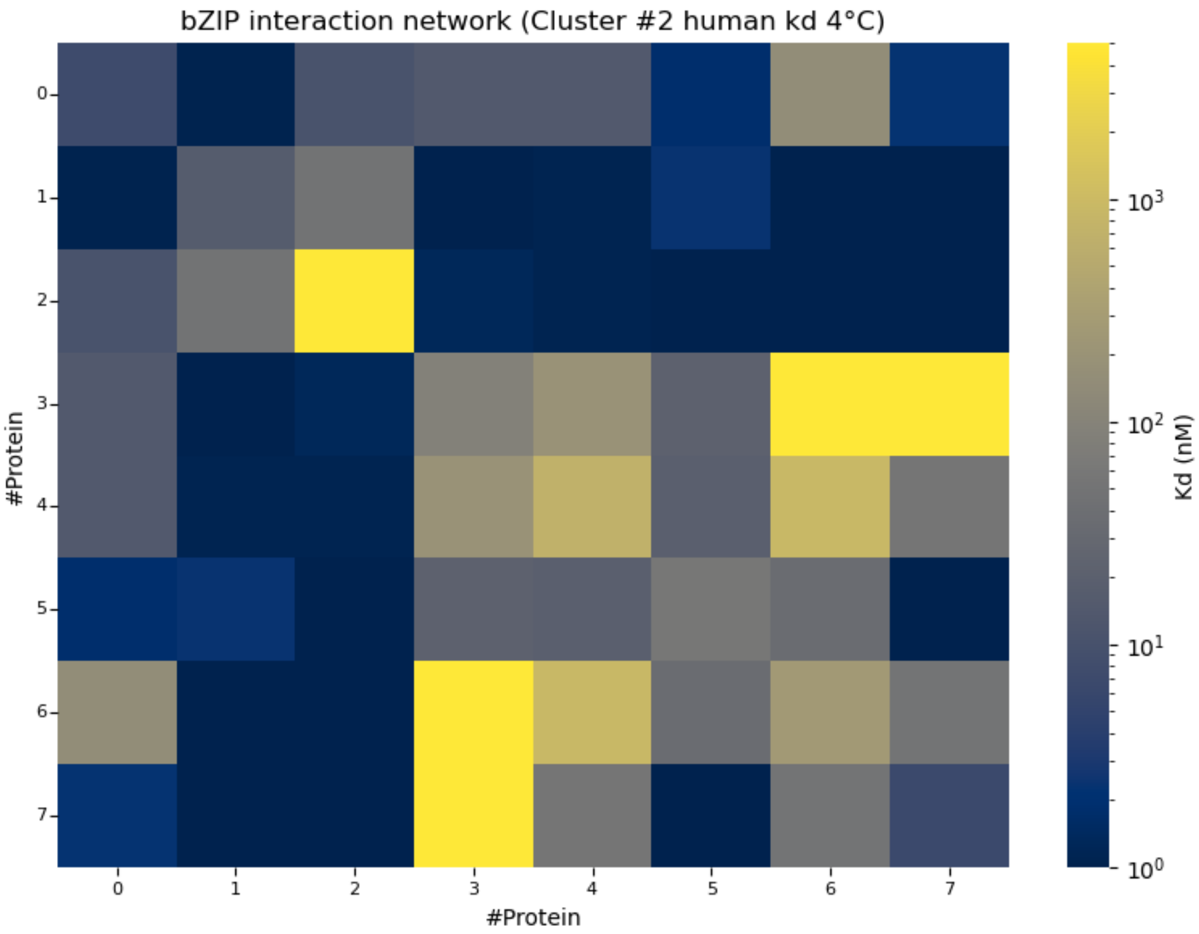


**Figure S17. Binding affinity matrix of *Homo sapiens* bZIP cluster #2 at 4°C.**

Cluster #2 comprises eight bZIP proteins connected by 33 interactions among 36 possible unique dimer pairs. Its connectivity density of 91.7% makes it the most densely connected of the selected Homo sapiens clusters at 4°C. The matrix presents dissociation constants $K_d$ $(nM)$. Protein indices start at 0 and follow the order listed for this cluster in **Table S3**. JUN and ATF3, corresponding to indices 1 and 5, respectively, are the most highly connected proteins within the cluster.

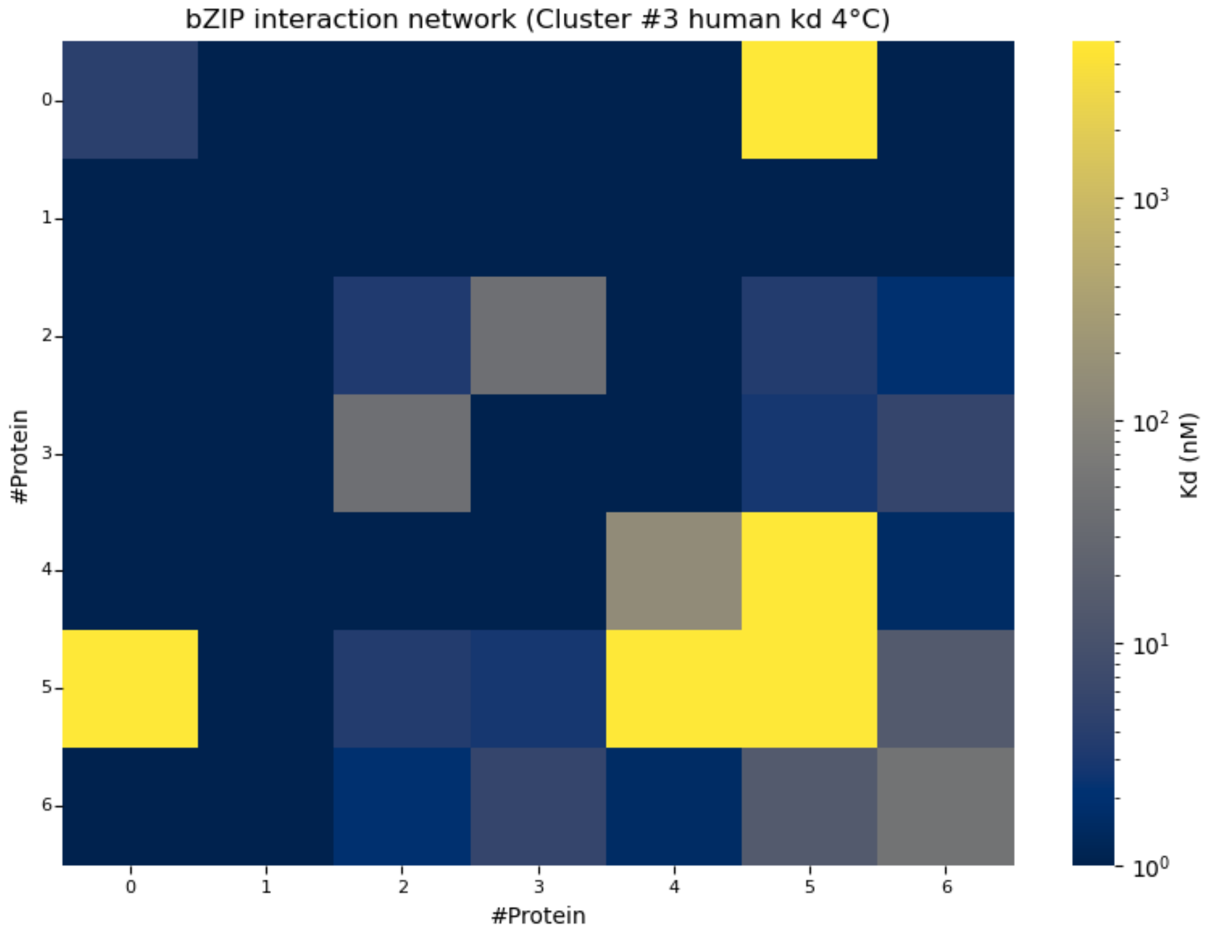


**Figure S18. Binding affinity matrix of *Homo sapiens* bZIP cluster #3 at 4°C.**

The matrix presents dissociation constants $K_d$ $(nM)$ from Cluster #3. This cluster comprises seven bZIP proteins, with interactions present in 25 of the 28 possible unique dimer pairs. Only three possible pairs lack an interaction in the curated network, giving this cluster a connectivity density of 89.3%, the second highest among the selected *Homo sapiens* clusters at 4°C. Protein indices start at 0 and follow the order listed for this cluster in **Table S3**. CEBPG and CEBPE, indexed as 1 and 3, respectively, are the most highly connected proteins, linking them to multiple potential dimerization partners.

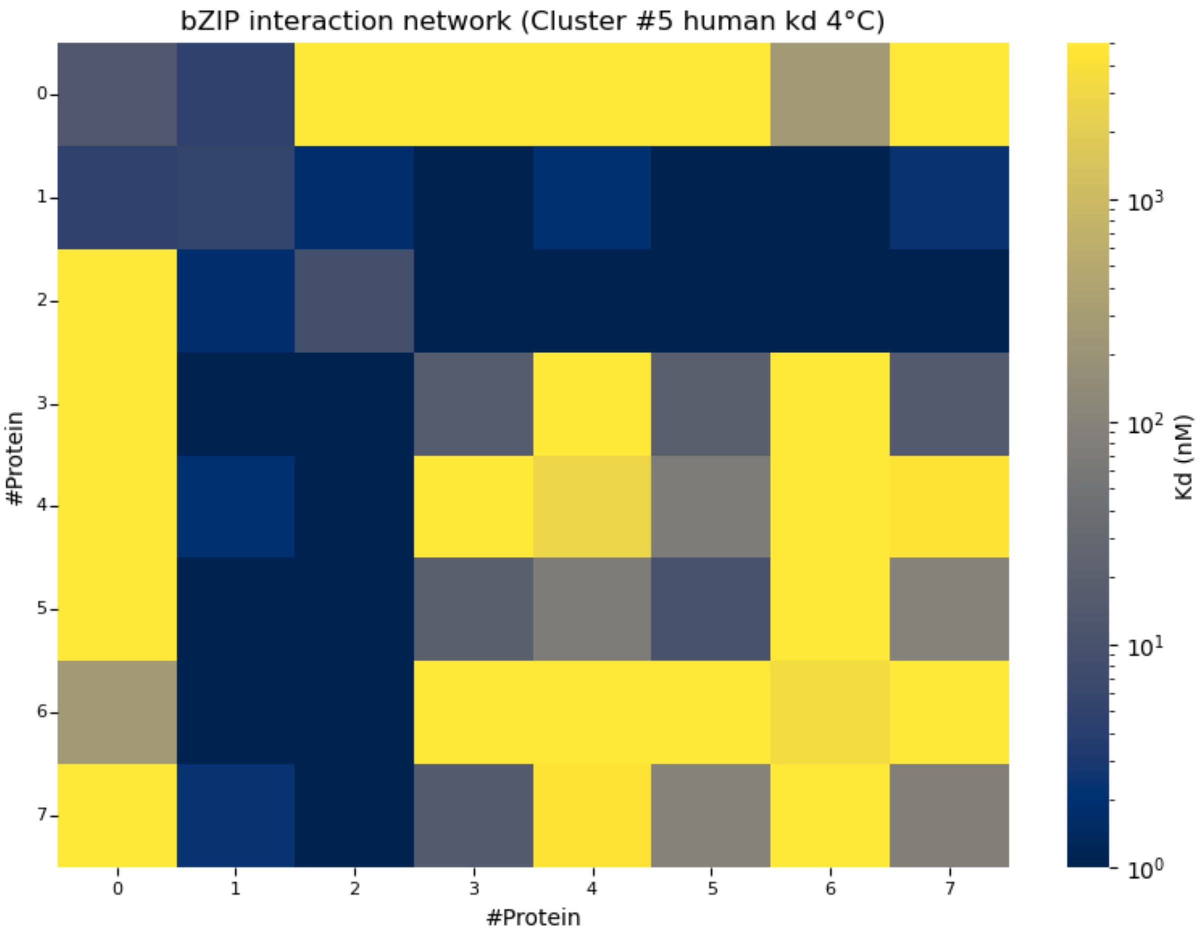


**Figure S19. Binding affinity matrix of *Homo sapiens* bZIP cluster #5 at 4°C.**

This matrix presents dissociation constants $K_d$ $(nM)$ from Cluster #5. This cluster contains eight bZIP proteins and 26 interactions among 36 possible unique dimer pairs, corresponding to a connectivity density of 72.2%. Although it has the same number of proteins as cluster #2 (**Figure S17**), it contains seven fewer interactions, providing a similarly sized network with lower internal connectivity. Protein indices start at 0 and follow the order listed for this cluster in **Table S3**. MAFF and MAFG, indexed as 1 and 2, respectively, are the most highly connected proteins.

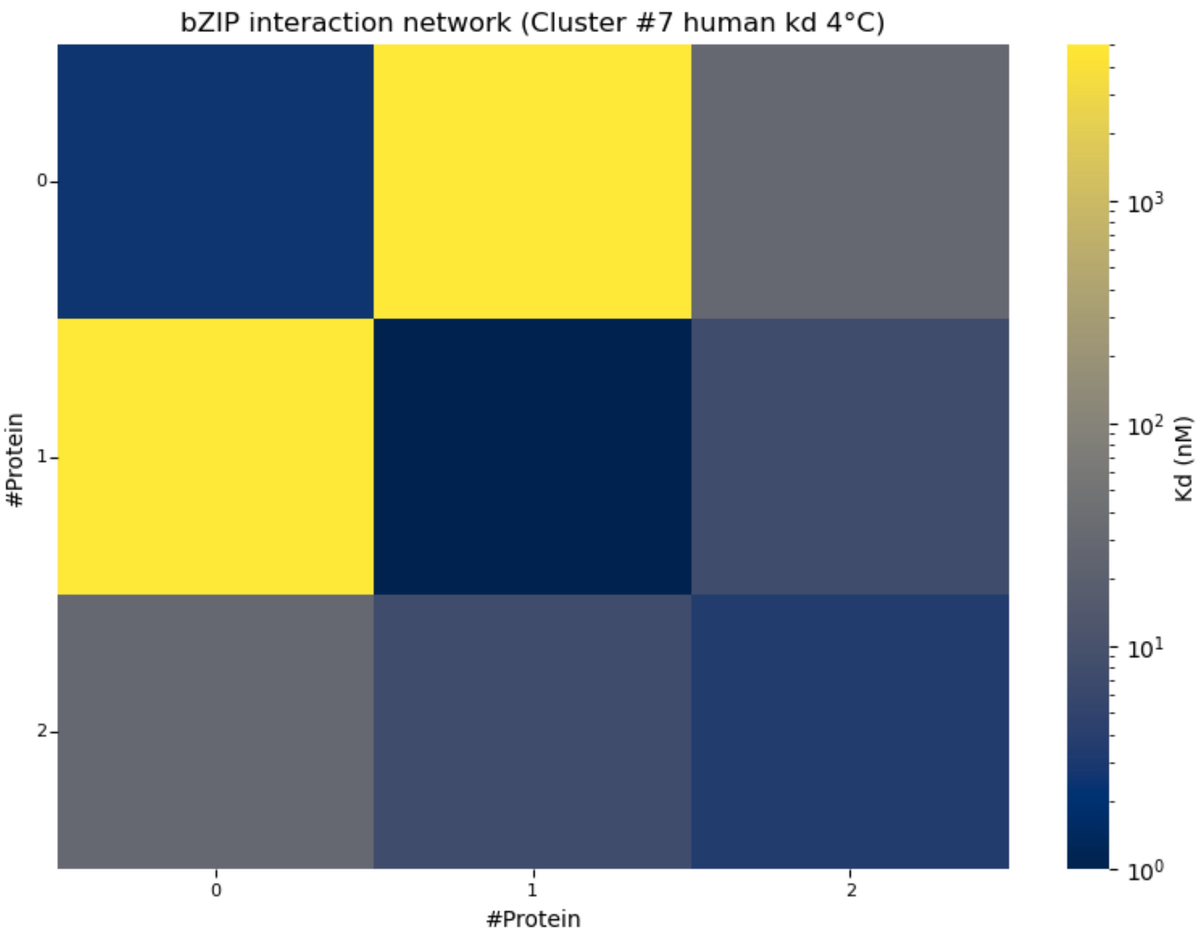


**Figure S20. Binding affinity matrix of *Homo sapiens* bZIP cluster #7 at 4°C.**

The matrix from Cluster #7 presents dissociation constants $K_d$ $(nM)$. It comprises three bZIP proteins and contains five of the six possible unique dimer interactions, yielding a connectivity density of 83.3%. This compact network accommodates two input monomers and one accessory monomer, with only one possible dimer pair lacking an interaction in the curated network. Protein indices start at 0 and follow the order listed for this cluster in **Table S3**. CREB3L3 and CREB3L1, corresponding to indices 1 and 2, respectively, have the highest connectivity.

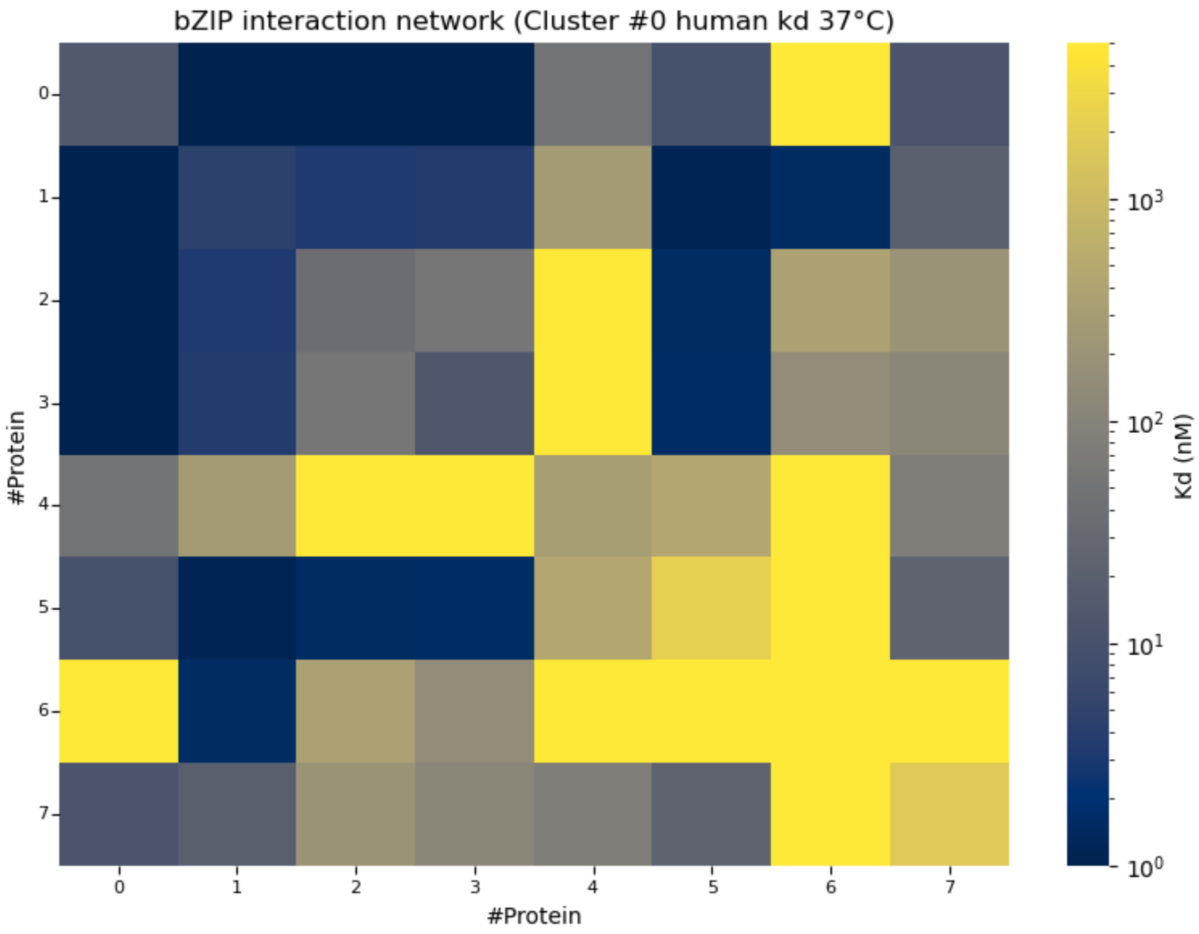


**Figure S21. Binding affinity matrix of *Homo sapiens* bZIP cluster #0 at 37°C.**

The matrix from Cluster #0 at 37°C presents dissociation constants $K_d$ $(nM)$ and comprises eight bZIP proteins connected by 29 interactions among 36 possible unique dimer pairs, corresponding to a connectivity density of 80.6%. This community shares several proteins with cluster #3 at 4°C (**Figure S18**), although its composition differs and its size increases from seven to eight proteins, which means more interactions overall but shows a smaller proportion of its possible dimer pairs. Protein indices start at 0 and follow the order listed for this cluster in **Table S4**.

#### Supplementary Note S2: Logic Functions Computed by bZIP Clusters

This section summarizes the logic functions successfully computed *in silico* by the selected bZIP clusters through optimization of accessory monomer concentrations while keeping experimental binding affinities fixed. **Table S5** reports performance across all 16 two-input Boolean functions, whereas **Table S6** presents results for the proposed benchmark suite of 17 three-input functions. Together, these tables identify the functions achieved by each evaluated configuration and summarize its computational versatility.

**Table S5. Two-input Boolean functions computed by the selected bZIP clusters.**

The table summarizes the two-input Boolean functions computed by the selected bZIP clusters. Each row corresponds to a bZIP cluster, identified by organism and cluster number, and each function column represents one of the 16 possible two-input Boolean functions. Four-digit binary sequences specify the target outputs for input states ordered as 00, 01, 10, and 11. Green entries indicate functions successfully computed under the classification criteria described in Methods, whereas yellow entries indicate functions not successfully computed. The final column reports computational versatility as the number of functions achieved out of 16. Each successful function corresponds to an optimized configuration of accessory monomer concentrations and a selected output dimer within the same fixed-affinity cluster.

| **Organism** | **Cluster** | **FALSE** | **AND** | **A AND NOT B** | **BUFFER A** | **NOT A AND B** | **BUFFER B** | **XOR** | **OR** | **NOR** | **XNOR** | **NOT B** | **B IMPLIES A** | **NOT A** | **A IMPLIES B** | **NAND** | **TRUE** | **Versatility** |
|---|---|---|---|---|---|---|---|---|---|---|---|---|---|---|---|---|---|---|
| | | 0000 | 0001 | 0010 | 0011 | 0100 | 0101 | 0110 | 0111 | 1000 | 1001 | 1010 | 1011 | 1100 | 1101 | 1110 | 1111 | |
| ***N. V.*** | #0 | | | | | | | | | | | | | | | | | 14/16 |
| ***N. V.*** | #3 | | | | | | | | | | | | | | | | | 15/16 |
| ***N. V.*** | #5 | | | | | | | | | | | | | | | | | 12/16 |
| ***C. I.*** | #0 | | | | | | | | | | | | | | | | | 5/16 |
| ***C. I.*** | #1 | | | | | | | | | | | | | | | | | 7/16 |
| ***C. I.*** | #5 | | | | | | | | | | | | | | | | | 12/16 |
| ***C. I.*** | #7 | | | | | | | | | | | | | | | | | 15/16 |
| ***H. S.*** | #0 | | | | | | | | | | | | | | | | | 11/16 |
| ***H. S.*** | #2 | | | | | | | | | | | | | | | | | 14/16 |
| ***H. S.*** | #3 | | | | | | | | | | | | | | | | | 14/16 |
| ***H. S.*** | #5 | | | | | | | | | | | | | | | | | 11/16 |
| ***H. S.*** | #7 | | | | | | | | | | | | | | | | | 5/16 |

**Table S6. Three-input Boolean functions computed by the selected bZIP clusters.**

The table summarizes the three-input Boolean functions computed by the selected bZIP clusters across the proposed 17-function benchmark suite. Each row identifies the organism and cluster, while each function column represents a target Boolean operation. Eight-digit binary sequences specify the target outputs for input states ordered as 000, 001, 010, 011, 100, 101, 110, and 111. Green entries indicate functions successfully computed under the classification criteria described in Methods, whereas yellow entries indicate functions not successfully computed. The final column reports computational versatility as the number of functions achieved out of the 17 evaluated. The three-input combinations were NV11, NV35, and NV34 for the *N. vectensis* cluster #0 classified as poorly connected; NV3, NV7, and NV30 for the *N. vectensis* cluster #3 classified as intermediately connected; and CI1, CI18, and CI31 for the *C. intestinalis* cluster #7 classified as intermediately connected. Each successful function corresponds to an optimized configuration of accessory monomer concentrations and a selected output dimer within the same fixed-affinity cluster.

| **Organism / Cluster** | **AND** 00000001 | **OR** 01111111 | **MAJ** 00010111 | **XOR** 01101001 | **MUX** 00011011 | **NAND** 11111110 | **NOR** 10000000 | **XNOR** 10010110 | **MIN** 11101000 | **EXACTLY ONE** 01101000 | **EXACTLY TWO** 00010110 | **BUFFER A** 00001111 | **BUFFER B** 00110011 | **BUFFER C** 01010101 | **NOT A** 11110000 | **NOT B** 11001100 | **NOT C** 10101010 | **Versatility** |
|---|---|---|---|---|---|---|---|---|---|---|---|---|---|---|---|---|---|---|
| ***N. V. #0*** | | | | | | | | | | | | | | | | | | 9/17 |
| ***N. V. #3*** | | | | | | | | | | | | | | | | | | 13/17 |
| ***C. I. #7*** | | | | | | | | | | | | | | | | | | 11/17 |

#### Supplementary Note S3: Logic Functions Computed by synthetic dimerization networks.

This section describes the search for minimal synthetic dimerization networks capable of computing two-input Boolean functions, following the optimization framework presented in our previous work[2]. Each function was evaluated by jointly optimizing binding affinities and accessory monomer concentrations. The search began with three monomer networks, comprising two inputs and one accessory, and was extended to four monomers to obtain the remaining functions. Each successful configuration defines a distinct synthetic network with its own optimized affinities and concentrations.

##### S3.1 Search for Minimal Synthetic Networks Using Genetic Algorithm Optimization

Following the optimization procedure described in our previous work[2], we used a genetic algorithm to identify minimal synthetic dimerization networks capable of computing two-input Boolean functions. The search began with three monomers, comprising two inputs and one accessory monomer. Each candidate solution encoded the pairwise binding affinities and accessory monomer concentration, which were initialized randomly and jointly optimized.

For each candidate, EQTK calculated equilibrium dimer concentrations across the four input combinations of the target truth table. The concentration profile of each candidate output dimer was compared with the desired response using a sum-of-squared-errors objective, which was converted into a normalized fitness score ranging from zero to one. A fitness threshold of 0.90 was used as the acceptance criterion, and all selected configurations met or exceeded this threshold.

Optimization was performed over 400 generations with a population of 360 individuals, binary tournament selection, and a dispersion parameter of 3. An elitism count of one preserved the highest-fitness individual across generations.

The three-monomer search yielded configurations for 14 of the 16 target functions. Since no configurations meeting the fitness criterion were identified for XOR or OR, the network was expanded to four monomers and the optimization was repeated with two accessory components. Successful configurations were then obtained for both remaining functions. Each selected network was defined by its optimized binding affinities, accessory concentrations, and output dimer.

**Table S7. Two-input Boolean functions obtained through joint optimization of binding affinities and accessory concentrations in three- and four-monomer synthetic networks.**

The table summarizes the two-input Boolean functions obtained with three- and four-monomer synthetic dimerization networks. Four-digit binary sequences indicate the target outputs. Dark green identifies the 14 functions achieved with three monomers, whereas light green identifies XOR and OR, which were achieved with networks of four monomers. Binding affinities and accessory monomer concentrations were optimized independently for each function. A fitness threshold of 0.90 was used as the acceptance criterion. Together, these configurations encompass all 16 two-input Boolean functions.

| Number of monomers | FALSE | AND | A AND NOT B | BUFFER A | NOT A AND B | BUFFER B | XOR | OR | NOR | XNOR | NOT B | B IMPLIES A | NOT A | A IMPLIES B | NAND | TRUE | Versatility |
|---|---|---|---|---|---|---|---|---|---|---|---|---|---|---|---|---|---|
| | 0000 | 0001 | 0010 | 0011 | 0100 | 0101 | 0110 | 0111 | 1000 | 1001 | 1010 | 1011 | 1100 | 1101 | 1110 | 1111 | |
| 3 | | | | | | | | | | | | | | | | | 16/16 |
| 4 | | | | | | | | | | | | | | | | | |

#### Supplementary Note S4: Influence of Input Monomer Selection on Computational Versatility

This section examines whether selecting different proteins as inputs changes the number of logic functions that a cluster can compute. Using *Nematostella vectensis* cluster #0 at 4 °C, we tested all 21 possible pairs among its seven proteins. For each pair, the two selected proteins served as inputs, while the concentrations of the remaining five accessory proteins were optimized separately for each of the 16 two-input Boolean functions. The proteins comprising the cluster and their binding affinities remained unchanged throughout the analysis. **Table S8** reports the number of functions achieved by each input pair alongside its connectivity, showing how computational versatility varies with input selection.

**Table S8. Input-pair selection and computational versatility of *Nematostella vectensis* bZIP cluster #0 at 4°C.**

The table compares computational versatility across all 21 input pairs in a cluster containing seven proteins and 20 interactions. Each row identifies the two input proteins and their individual numbers of connections within the cluster. Total input connections represent the combined number of unique interactions involving either input, with shared interactions counted only once. Computed gates indicate the number of two-input Boolean functions successfully achieved, while versatility expresses this number as a fraction of the 16 functions evaluated. Performance ranges from 10 to 15 functions depending on the input pair. The most connected pair, NV31–NV23, achieves 12 functions, whereas NV11–NV10 achieves 15 despite having fewer combined connections, illustrating that greater input connectivity does not necessarily yield greater computational versatility.

| ***N. Vectensis* 4°C - #0** | | | | | | |
|---|---|---|---|---|---|---|
| **Cluster size:** | | | 7 monomers | | | |
| **Total cluster interactions:** | | | 20 dimers | | | |
| **Input 1** | **Input 1 connections** | **Input 2** | **Input 2 connections** | **Total inputs connections** | **Computed gates** | **Versatility** |
| NV11 | 3 | NV5 | 6 | 8 | 14 | 0.8750 |
| NV11 | 3 | NV10 | 5 | 8 | 15 | 0.9375 |
| NV11 | 3 | NV31 | 7 | 9 | 14 | 0.8750 |
| NV11 | 3 | NV23 | 7 | 9 | 14 | 0.8750 |
| NV11 | 3 | NV35 | 5 | 8 | 14 | 0.8750 |
| NV11 | 3 | NV34 | 2 | 5 | 13 | 0.8125 |
| NV5 | 6 | NV10 | 5 | 10 | 13 | 0.8125 |
| NV5 | 6 | NV31 | 7 | 12 | 14 | 0.8750 |
| NV5 | 6 | NV23 | 7 | 12 | 14 | 0.8750 |
| NV5 | 6 | NV35 | 5 | 10 | 14 | 0.8750 |
| NV5 | 6 | NV34 | 2 | 8 | 13 | 0.8125 |
| NV10 | 5 | NV31 | 7 | 11 | 14 | 0.8750 |
| NV10 | 5 | NV23 | 7 | 11 | 14 | 0.8750 |
| NV10 | 5 | NV35 | 5 | 9 | 15 | 0.9375 |
| NV10 | 5 | NV34 | 2 | 7 | 10 | 0.6250 |
| NV31 | 7 | NV23 | 7 | 13 | 12 | 0.7500 |
| NV31 | 7 | NV35 | 5 | 11 | 13 | 0.8125 |
| NV31 | 7 | NV34 | 2 | 8 | 13 | 0.8125 |
| NV23 | 7 | NV35 | 5 | 11 | 15 | 0.9375 |
| NV23 | 7 | NV34 | 2 | 8 | 13 | 0.8125 |
| NV35 | 5 | NV34 | 2 | 7 | 13 | 0.8125 |

### Supplementary Note S5: Mathematical Proofs and Thermodynamic Scaling Framework

#### S5.1 Non-Dimensionalization of Mass-Action Equilibrium

Consider a network of $N$ monomeric species $M_1, M_2, \dots, M_N$ with total loaded species concentrations $C_{i,\text{tot}}$ (representing the sum of species $i$ in both free monomeric and bound dimeric forms) forming dimeric complexes $D_{ij}$. Thermodynamic equilibrium is governed by mass action and mass conservation:

$$[D_{ij}]_{\text{eq}} = K_{ij}[M_i]_{\text{eq}}[M_j]_{\text{eq}} \qquad \text{Eq. S1}$$

$$C_{i,\text{tot}} = [M_i]_{\text{eq}} + \sum_{j=1}^{N}(1+\delta_{ij})[D_{ij}]_{\text{eq}} \qquad \text{Eq. S2}$$

Where $\delta_{ij} = 1$ if $i = j$ (homodimer) and $\delta_{ij} = 0$ if $i \neq j$ (heterodimer). Defining the dimensionless free monomer fraction $x_i \in [0,1]$ as $x_i = \frac{[M_i]_{\text{eq}}}{C_{i,\text{tot}}}$ and substituting mass action into mass conservation yields:

$$C_{i,\text{tot}} = x_i C_{i,\text{tot}} + \sum_{j=1}^{N}(1+\delta_{ij})K_{ij}(x_i C_{i,\text{tot}})(x_j C_{j,\text{tot}}) \qquad \text{Eq. S3}$$

Dividing through by $C_{i,\text{tot}}$ introduces the dimensionless affinity product $\Gamma_{ij} = K_{ij}C_{j,\text{tot}}$:

$$1 = x_i\left(1 + \sum_{j=1}^{N}(1+\delta_{ij})\Gamma_{ij}\,x_j\right) \quad for \ = 1,2,3,\dots,N \qquad \text{Eq. S4}$$

**Eq. S4** defines a system of non-linear algebraic equations $\boldsymbol{F}(\boldsymbol{x},\Gamma) = \boldsymbol{0}$. Because the governing equations depend strictly on the dimensionless matrix $\Gamma$, the vector of free monomer fractions $\boldsymbol{x}$ is mathematically invariant under any parameter transformation that preserves $\Gamma$.

#### S5.2 Proof of Scenario A: Exact Fractional Invariance ($\boldsymbol{K'} = \frac{\boldsymbol{K}}{\boldsymbol{\alpha}}$)

If total loaded concentrations are scaled by $\alpha$ ($C'_{\text{tot}} = \alpha C_{\text{tot}}$) and affinities are scaled as $K'_{ij} = \frac{K_{ij}}{\alpha}$, the new dimensionless parameter matrix $\Gamma'$ satisfies:

$$\Gamma'_{ij} = K'_{ij} C'_{j,\text{tot}} = \left(\frac{K_{ij}}{\alpha}\right)\left(\alpha C_{j,\text{tot}}\right) = K_{ij} C_{j,\text{tot}} = \Gamma_{ij} \qquad \text{Eq. S5}$$

Since $\Gamma' = \Gamma$, the solution vector to **Eq. S4** is unchanged ($x'_i = x_i$). The fraction of total system mass converted into dimer $D_{ij}$, defined as $f_{D_{ij}} = \frac{[D_{ij}]_{\text{eq}}}{C_{\text{total}}}$, transforms as:

$$f'_{D_{ij}} = \frac{K'_{ij}[M_i]'[M_j]'}{C'_{\text{total}}} = \frac{\left(\frac{K_{ij}}{\alpha}\right)\left(x_i \alpha C_{i,\text{tot}}\right)\left(x_j \alpha C_{j,\text{tot}}\right)}{\alpha C_{\text{total}}} = \frac{K_{ij} C_{i,\text{tot}} C_{j,\text{tot}} x_i x_j}{C_{\text{total}}} = f_{D_{ij}} \qquad \text{Eq. S6}$$

Thus, fractional species distributions, relative equilibrium ratios, and binary logic state margins remain 100% identical across concentration scales.

#### S5.3 Proof of Scenario B: Quadratic Scaling in Dominant Monomer Regimes

In experiments where natural affinities remain unadjusted ($K'_{ij} = K_{ij}$), uniform concentration scaling $C'_{\text{tot}} = \alpha C_{\text{tot}}$ modifies species abundances. In the asymptotic limit of dominant monomeric species/low conversion ($x_i \approx 1$, where $[M_i]_{\text{eq}} \approx C_{i,\text{tot}}$), dimer output is given by:

$$\left[D_{ij}\right]_{\text{eq}} = K_{ij}[M_i]_{\text{eq}}\left[M_j\right]_{\text{eq}} \approx K_{ij} C_{i,\text{tot}} C_{j,\text{tot}} \qquad \text{Eq. S7}$$

Under uniform scaling $C'_{\text{tot}} = \alpha C_{\text{tot}}$:

$$\left[D_{ij}\right]'_{eq} = K_{ij}\left(\alpha C_{i,tot}\right)\left(\alpha C_{j,tot}\right)\left(x'_i x'_j\right) \approx \alpha^2 K_{ij} C_{i,tot} C_{j,tot}\left(x_i x_j\right) = \alpha^2\left[D_{ij}\right]_{eq} \qquad \text{Eq. S8}$$

This quadratic relation ($\approx \alpha^2$) holds uniformly across all network interactions, including reporter output dimers formed by input–input, input–accessory, or accessory–accessory pairings, provided accessory baseline abundances are scaled proportionally alongside the input vector.

#### S5.4 Departure from Asymptotic Limit and Transition to Saturation ($\alpha$)

Evaluating this framework within our experimental dataset ($K_a \in [2 \times 10^{-4}, 1]$ $nM^{-1}$, $C \in [10^{-3}, 10^{2}]$ $nM$) identifies two operational regimes for target reporter outputs:

1. **Unsaturated / Low Output Regime ($\Gamma_{ij} \ll 1$):** In basal background output regimes, weak binding pathways ($K_a \approx 2 \times 10^{-4}$ $nM^{-1}$), or low concentration states ($C \approx 10^{-3}$ $nM$), $\Gamma_{ij} \ll 1$. Free monomer depletion is negligible ($x_i \approx 1$), and background dimer formation scales quadratically ($\approx \alpha^2$), remaining near zero in absolute concentration.

2. **Saturated / High Output Regime ($\Gamma_{ij} \gg 1$):** In high output signal regimes governed by high affinities ($K_a \approx 1$ nM$^{-1}$) and high monomer loading ($C \approx 10^2$ nM), $\Gamma_{ij} \gg 1$. Significant monomer depletion occurs ($x_i < 1$). In this limit, target output dimer formation is constrained by reactant stoichiometry rather than collision frequency:

$$\lim_{\Gamma\to\infty}\left[D_{ij}\right]_{\mathrm{eq}} \approx \min\left(C_{i,\mathrm{tot}}, C_{j,\mathrm{tot}}\right) \quad \text{Eq. S9}$$

Scaling concentrations by yields a near-linear output response:

$$\lim_{\Gamma\to\infty}\left[D_{ij}\right]'_{\mathrm{eq}} \approx \min\left(\alpha C_{i,\mathrm{tot}}, \alpha C_{j,\mathrm{tot}}\right) = \alpha \cdot \min\left(C_{i,\mathrm{tot}}, C_{j,\mathrm{tot}}\right) \approx \alpha\left[D_{ij}\right]_{\mathrm{eq}} \quad \text{Eq. S10}$$

#### Impact on Logic Signal Margin ($\Delta\left[D_{\mathrm{output}}\right]$)

Because low-output baseline noise scales quadratically from near-zero levels while high-output signal scales robustly, scaling concentrations to micromolar cell-free regimes expands the absolute signal window ($\Delta\left[D_{\mathrm{output}}\right] = \left[D_{\mathrm{high}}\right]' - [D_{\mathrm{low}}]'$), improving signal-to-noise ratio and digital contrast for experimental readout.
